# Machine learning pattern recognition and differential network analysis of gastric microbiome in the presence of proton pump inhibitor treatment or *Helicobacter pylori* infection

**DOI:** 10.1101/2020.03.24.005587

**Authors:** Sara Ciucci, Claudio Durán, Alessandra Palladini, Umer Z. Ijaz, Francesco Paroni Sterbini, Luca Masucci, Giovanni Cammarota, Gianluca Ianiro, Pirjo Spuul, Michael Schroeder, Stephan W. Grill, Bryony N. Parsons, D. Mark Pritchard, Brunella Posteraro, Maurizio Sanguinetti, Giovanni Gasbarrini, Antonio Gasbarrini, Carlo Vittorio Cannistraci

## Abstract

Although long thought to be a sterile and inhospitable environment, the stomach is inhabited by diverse microbial communities, co-existing in a dynamic balance. Long-term use of orally administered drugs such as Proton Pump Inhibitors (PPIs), or bacterial infection such as *Helicobacter pylori*, cause significant microbial alterations. Yet, studies revealing how the commensal bacteria re-organize, due to these perturbations of the gastric environment, are in the early phase. They mainly focus on the most prevalent taxa and rely on linear techniques for multivariate analysis.

Here we disclose the importance of complementing linear dimensionality reduction techniques such as Principal Component Analysis and Multidimensional Scaling with nonlinear approaches derived from the physics of complex systems. Then, we show the importance to complete multivariate pattern analysis with differential network analysis, to reveal mechanisms of re-organizations which emerge from combinatorial microbial variations induced by a medical treatment (PPIs) or an infectious state (*H. pylori*).

## Introduction

The gastric environment with its microbiota is the active gate that regulates access to the whole gastrointestinal tract, and therefore it has a remarkable impact on the correct functionality of the entire human organism. Recent studies have revealed that many orally administered drugs can perturb the elegant balance of the gastric flora ^1,2^. However, not all of them cause permanent adverse effects and particular attention should be addressed to drugs that are frequently prescribed and administered for long periods. They can cause permanent unbalance of the gastric microbiota that might generate adverse side effects for the patient’s health. Since the introduction of proton pump inhibitors (PPIs) into clinical practice more than 25 years ago, PPIs have become the mainstay in the treatment of gastric-acid-related diseases ^3^. PPIs are potent agents that block acid secretion by gastric parietal cells by binding covalently to and inhibiting the hydrogen/potassium (H^+^/K^+^)-ATPases (or proton pumps), and additionally they can bind non-gastric H^+^/K^+^-ATPases, both on human cells and on bacteria and fungi, such as *Helicobacter pylori* (*H. pylori*)^4–6^.

PPIs are drugs of first choice for peptic ulcers (PU) and their complications (e.g. bleeding), gastroesophageal reflux disease (GERD), nonsteroidal anti-inflammatory drug (NSAID)-induced gastrointestinal (GI) lesions, Zollinger-Ellison syndrome and dyspepsia ^3,7,8^. In particular, dyspepsia is a common clinical problem characterized by symptoms (e.g. epigastric pain, burning, postprandial fullness, or early satiation) originating from the gastroduodenal region ^9^. The potent gastric-acid suppression drugs PPIs can treat the most frequent causes of dyspepsia including GERD, medication-induced gastritis, and peptic ulcers, thus minimizing the need for costly and invasive testing, and moreover are currently recommended to eradicate *H. pylori* infection, in combination to antibiotics ^7,9,10^. Nevertheless, some patients are resistant or partial responders to empiric PPI therapy, and continue to have dyspepsia ^7^.

Additionally, there is growing evidence that these medications are associated with increased rates of pharyngitis and upper and lower respiratory tract infections ^11^. Their long-term overutilization has been associated with potential adverse effects. For instance: the development of corpus predominant atrophic gastritis in *H. pylori* positive patients (that is a precursor of gastric cancer), enteric infections (especially *Clostridium difficile*-associated diarrhoea), increased risk of fundic gland polyps, hypomagnesaemia and hypocalcaemia, osteoporosis and bone fractures, vitamin and mineral deficiency, pneumonia, acute interstitial nephritis, and increased risk of drug–drug interactions, among others ^7,12–15^.

Consumption of such acid-suppressive medications has also been associated with changes in microbial composition and function of gut microbiota. More recent studies relying on amplicon-based metagenomic approaches, have shown that PPIs exert an effect on gastric, oropharyngeal, and lung microflora in children with a chronic cough ^11^, and have a significant impact on the gut microbiome in healthy subjects, with an increase of oral and pharyngeal bacteria and potential pathogenic bacteria ^16,17^. Furthermore, another study by Tsuda *et al.* ^18^ revealed that PPIs influence the bacterial composition of saliva, gastric fluid and stool in a cohort of adult dyspeptic patients. However, this latter study highlights how the influence of PPI administration on the fecal and gastric luminal microbiota is still controversial and further investigation is required to understand the interaction between PPIs and non-*H. pylori* bacteria. Hence, this represents the first reason that motivates the present study.

In fact, by irreversibly blocking H^+^/K^+^-ATPases, PPIs inhibit gastric acid secretion by gastric parietal cells, which results in a higher intragastric pH, meaning the microenvironment of this niche changes, hence allowing more bacteria to survive the gastric acid barrier ^4,5,16^. The use of PPIs and higher gastric pH were indeed correlated with the overgrowth of non-*H. pylori* bacterial flora in the stomach of patients with gastric-reflux and PPIs were shown to aggravate gastritis because of co-infection with *H. pylori* and non–*H. pylori* bacterial species ^4,14,19,20^. However, PPIs may also affect the gastrointestinal microbiome through pH-independent mechanisms, by directly targeting the proton pumps of naturally occurring bacteria by binding P-type ATPases (e.g. *H. pylori)* ^4,6^.

Attempts to detect patterns of PPI related gastrointestinal changes have been made in different studies ^21,22^ through linear multidimensional analysis techniques, such as Principal Component Analysis (PCA) and Multidimensional Scaling (MDS), also called Principal Coordinates Analysis (PCoA). Nevertheless, they failed to detect the effect of PPIs on gastric *fluid* samples ^21^, nor any significant PPI-related modification in esophageal ^21^ and gastric ^22^ *tissue* samples. This represents the second reason that motivates our investigation. Are these controversial results due to complex patterns that cannot be detected using linear analysis?

In this study, we show an unprecedented result: unlike linear approaches, Minimum Curvilinear Embedding (MCE) ^23^, which is a technique for *nonlinear* dimension reduction, discriminated both the esophageal and the gastric tissue microbial profiles of patients taking PPI medications from untreated ones when re-analyzing the data published in the abovementioned studies. This finding demonstrates the importance of routinely integrating the use of nonlinear multidimensional techniques into clinical metagenomic studies, since addressing nonlinearity could significantly modify the results and conclusions. Indeed, the absence of separation by means of linear transformations does not imply absence of separation in general, and nonlinear techniques could prove it, especially in complex datasets such as the ones generated in metagenomics 16S rRNA. As a matter of fact, the high throughput profiling of bacteria is frequently used in clinical studies, thus posing a challenge to efficient information retrieval: understanding how microbial community structure affects health and disease can indeed contribute to better diagnosis, prevention, and treatment of human pathologies ^24^.

The common practice in unsupervised dimension reduction data analysis is to consider only the first two (or three, less used) dimensions of mapping, and the goal is to visually explore the distribution of the samples and the incidence of significant patterns ^25^. This procedure is particularly useful in case of studies with small size datasets ^23^, to obtain unbiased (the labels are not used) confirmation of the separation between groups of samples for which diversity is theorized or expected.

Here, we will specifically analyse the many aforementioned 16S rRNA amplicons datasets to address the following pattern recognition questions: (1) Is PPI treatment affecting change on the microbiota of esophageal and gastric tissues in dyspeptic patients, regardless of the initial pathological infection due to *H. pylori*? (2) Is this PPI-induced change so dominant as to result in a discernible pattern in the first two dimensions of mapping by unsupervised dimension reduction? (3) Are linear techniques sufficient to bring out patterns in complex microbial data? Furthermore, using differential network analysis we will address from the systems point of view these other questions: (4) How is PPI affecting the microbiota in the gastric environment in dyspeptic patients? (5) What is the effect of *H. pylori* infection on gastric mucosal microflora? Both factors (PPI treatment and *H. pylori* infection) can influence the composition of the gastric microbiota, and this further analysis will help to understand the general (overall) behaviour of the microbial ecosystem under these conditions. Ultimately, this means that we will try to clarify and visualize via network representation how the bacterial cooperative organization is systemically altered either by the use of this acid suppressant drug in the gastric environment under dyspepsia, or by *H. pylori* infection in the gastric mucosa.

## Methods

### Dataset description

#### Amir3 (esophageal mucosa)

The 16S rRNA gene sequences were generated by Amir and colleagues ^21^ and are publicly available via the MG RAST database (http://metagenomics.anl.gov/linkin.cgi?project=5767). The dataset was obtained from 16 esophageal mucosal biopsies of eight individuals before and after eight weeks of PPI treatment. Two patients with heartburn presented normal oesophagogastroduodenoscopy (H) indicating that they present healthy oesophageal tissues but are exposed to gastric refluxate, four patients had oesophagitis (ES) and two had Barrett’s oesophagus (BE). Metagenomes were obtained by pyrosequencing 16S rRNA amplicons on the GS FLX system (Roche). Data were processed by replicating the bioinformatics workflow followed by Amir and colleagues ^21^ in order to obtain the matrix of the bacterial absolute abundance: sequence reads were analysed with the pipeline Quantitative Insights into Microbial Ecology (QIIME) v. 1.6.0 ^26^ using default parameters (sequences were removed if shorter than 200 nt, if they contained ambiguous bases or uncorrectable barcodes, or if the primer was missing). Operational Taxonomic Units (OTUs), that are clusters of sequences showing a pairwise similarity no lesser than 97%, were identified using the UCLUST algorithm (http://www.drive5.com/usearch/). The most abundant sequence in each cluster was chosen as the representative of its OTU, and this representative set of sequences was then used for taxonomy assignment by means of the Bayesian Ribosomal Database Project classifier ^27^ and aligned with PyNAST103. Chimeras, that are PCR artefacts, were identified using ChimeraSlayer ^28^ and removed. The Greengenes database, which was used for the annotation of the reads, additionally identifies groups of bacteria that are supported by whole genome phylogeny, but are not yet officially recognized by the Bergeys taxonomy, which is the reference taxonomy and is based on physiochemical and morphological traits. This results in a special annotation for some taxa, like *Prevotella,* that thus appears both with the general annotation, that is *Prevotella*, and with the special annotation, that is between square brackets, [*Prevotella*].

#### Amir4 (gastric fluid)

The dataset was generated by Amir and colleagues ^21^, and is public and available in the MG RAST database (http://metagenomics.anl.gov/linkin.cgi?project=5732). It comprises eight patients, whose gastric fluid was sampled at two different time points, that is before PPI treatment and after eight weeks of PPI treatment, for a total of 16 samples. The patients are the same described in Amir3. Metagenomes were obtained by pyrosequencing fragments of the 16S rRNA gene on the GS FLX system (Roche). Then the data were processed by replicating the same bioinformatics workflow followed by Amir and colleagues ^21^ that was described in the previous data description (Amir3), in order to obtain the matrix of the bacterial absolute abundance. As for Amir3, the Greengenes database was used for the annotation of the reads.

#### Paroni Sterbini (gastric mucosa)

The dataset was generated by Paroni Sterbini and colleagues ^22^, and is public and available in the NCBI Sequence Read Archive (SRA) (http://www.ncbi.nlm.nih.gov/sra, accession number SRP060417), where all details pertaining the sequencing experimental design are also reported. It contains 24 biopsy specimens of the gastric antrum from 24 individuals who were referred to the Department of Gastroenterology of Gemelli Hospital (Rome) with dyspepsia symptoms (i.e. heartburn, nausea, epigastric pain and discomfort, bloating, and regurgitation). Twelve of these individuals (PPI1 to PPI12) had been taking PPIs for at least 12 months, while the others (S1 to S12) were not being treated (naïve) or had stopped treatment at least 12 months before sample collection. In addition, 9 (5 treated and 4 untreated) were positive for *H. pylori* infection, where *H. pylori* positivity or negativity was determined by histology and rapid urease tests. Metagenomes were obtained by pyrosequencing fragments of the 16S rRNA gene on the GS Junior platform (454 Life Sciences, Roche Diagnostics). Then the sequence data were processed by replicating the bioinformatics workflow followed by Paroni Sterbini *et al.* ^22^, in order to obtain the matrix of the bacterial absolute abundance.

#### Parsons (gastric mucosa)

The dataset was generated by Parsons and colleagues ^29^, and is public and available in the EBI short-read archive (the European Nucleotide Archive, ENA) (https://www.ebi.ac.uk/ena, accession number PRJEB21104). In the original study, the authors focused on the analysis of gastric biopsy samples of 95 individuals (in groups representing normal stomach, PPI treated, *H. pylori-*induced gastritis, *H. pylori*-induced atrophic gastritis and autoimmune atrophic gastritis), selected from a larger prospectively recruited cohort patients who underwent diagnostic upper gastrointestinal endoscopy at Royal Liverpool University Hospital^29^. RNA extracted from gastric corpus biopsies was analysed using 16S rRNA sequencing (MiSeq). Then the sequence analysis was performed, as described by the authors in the supplementary methods of the original article ^29^. Here we focused on the analysis of gastric biopsy specimens (in total 42 samples) from normal stomach group (20 patients) and belonging to the *H. pylori* gastritis group (22 patients). As described in ^29^, patients in the normal stomach group showed normal endoscopy, no evidence of *H. pylori* infection by histology, rapid urease test or serology, were not treated by PPI and were normogastrinaemic. Patients in the *H. pylori* gastritis group were instead positive to *H. pylori* infection by urease test, histology and serology, were not taking PPI medication and were normogastrinaemic.

### Data exploration and visualization: the reason for unsupervised dimension reduction

The main reason to perform an unsupervised dimension reduction is to explore and visualize the most relevant sample patterns that should emerge in the first two dimensions of embedding (which represent the information of higher variability in the data) from the hidden multidimensional space of a dataset. The fact that the sample labels (if known) are not used for the data projection makes the analysis unsupervised. The advantage of performing an unsupervised analysis is both for data quality checking and to gather the main trends hidden in the data, independently from any hypothesis or knowledge available on the samples. This is particularly useful to discover the presence of interesting sub-groups inside the studied cohort or to detect the influence of confounding factors.

A final interesting advantage offered by unsupervised analysis is in small size datasets, where the number of samples *n* is significantly lower that the number of features *m*, a condition that unfortunately occurs in several metagenomic studies. When *n<<m* the application of supervised approaches can become problematic, because the supervised procedure of parameter learning can suffer from overfitting ^23,30,31^.

The mainstream multivariate methods to unsupervisedly explore data patterns in metagenomic studies are based on linear dimension reduction, in particular PCA ^32,33^ and MDS ^34,35^, also known as PCoA, methods that have been used to explore and visualize data structure in many metagenomic studies, from sponge ^36,37^ to gastric tissue microbiota ^22^. These tools perform a dimension reduction of the data either by *multidimensional variance analysis* (for instance PCA) or *dissimilarity embedding* (for instance MDS/PCoA). PCA collects uncorrelated variance in the multidimensional space, creating new synthetic orthogonal variables, which are linear combinations of the original ones, then plots the samples in a reduced space using the new variables that embody the largest orthogonal variances. MDS computes dissimilarities between every pair of samples, plotting the Euclidean part of these dissimilarities as distances between every pair of points (MDS) in a reduced space, in this way the linear part of the sample relations can be represented.

### The Tripartite-Swiss-Roll dataset

In order to test and visualize how the algorithms could detect nonlinearity, we performed the analyses on the Tripartite-Swiss-Roll dataset: an artificial dataset characterized by nonlinear structures and generated as discretization of the manifold associated to a Swiss-Roll function ^38^ in a three-dimensional (3D) space. Indeed, it is a synthetic dataset obtained as the partition in three sections of a discrete Swiss-Roll manifold depicted in a three-dimensional space ^38^. It reproduces the typical nonlinearity (given by the Swiss-Roll shape) and the discontinuity (given by the tripartition of the manifold), that we do not see and that are often hidden in the multidimensional representation of our samples. See the illustration in the original 3D-space of the Tripartite-Swiss-Roll dataset in Fig. 1A. This dataset is useful to introduce readers, not expert with nonlinear data analysis, to the basic concepts of nonlinear dimension reduction and therefore to facilitate their understanding of the new proposed methodologies for nonlinear dimension reduction.

**Figure 1.**
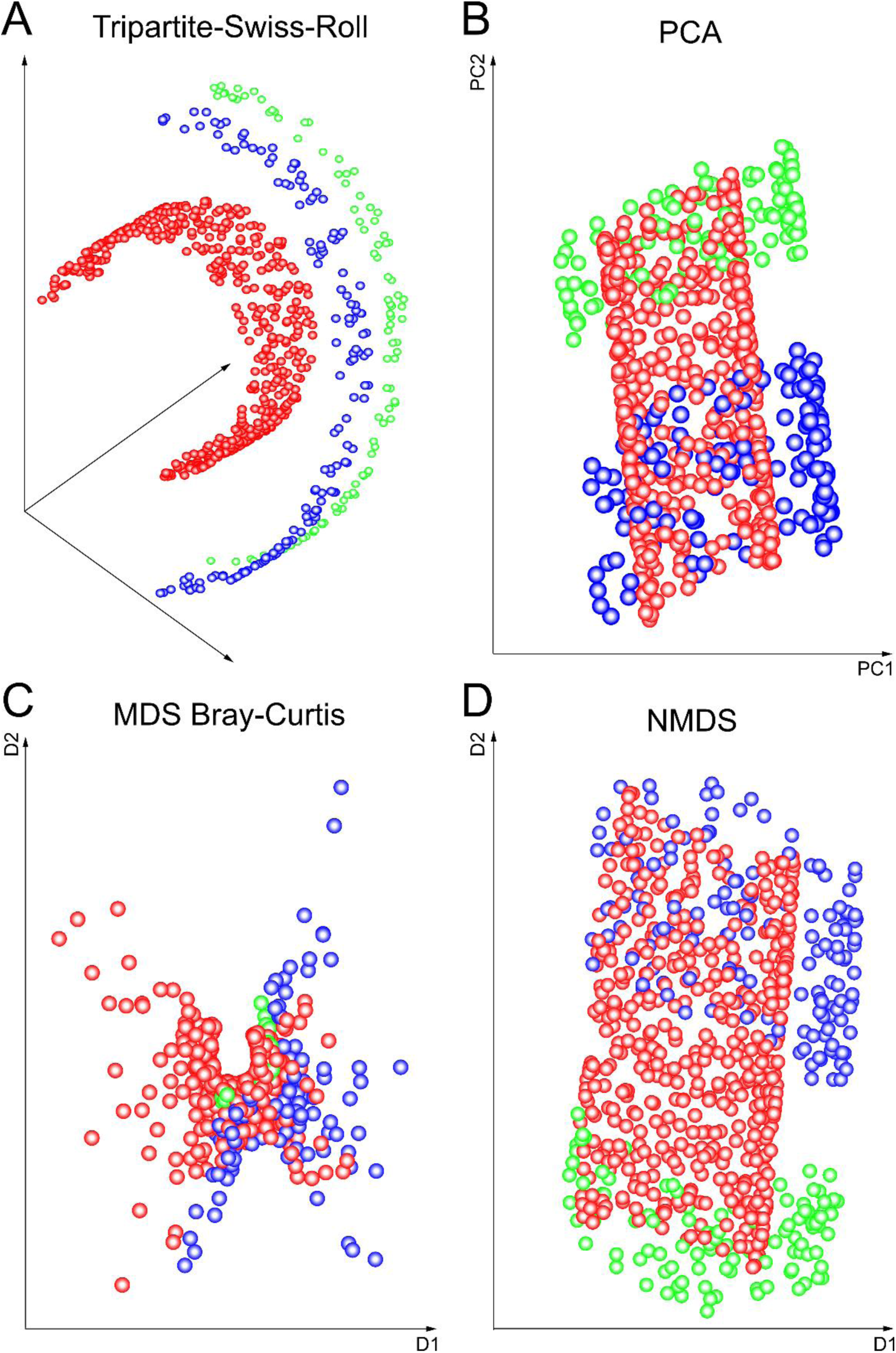
The Tripartite-Swiss-Roll as an example of data nonlinear organization. A) Tripartite-Swiss-Roll; B) PCA; C) MDS (Bray-Curtis dissimilarity); D) NMDS (Sammon Mapping). The three different colours (red, blue and green) represent the three partitions of the Swiss-roll manifold. This figure shows the inability of PCA, MDS and NMDS to reveal the inner nonlinear structure of the Tripartite-Swiss-Roll, which appears collapsed (B, D) or with a horseshoe shape (C).

### PCA, MDS (or PCoA) and LDA

Below, we report some of the PCA major advantages and drawbacks, that were pinpointed in a recent study on multidimensional population genomics ^39^, and of other conventional dimensional reduction techniques employed for the analysis of metagenomic data.

PCA is time-efficient, parameter-free and straightforward to interpret, yet it strives to resolve structure in datasets with few samples and highly numerous features, which enclose nonlinear patterns. Therefore, PCA can occasionally fail to reveal differences among samples, even when differences are known a-priori, which means it can also miss represent hidden nonlinear relations among the samples in the feature space. For instance, see the illustration of the PCA two-dimension reduction mapping of the Tripartite-Swiss-Roll dataset in Fig. 1B. PCA clearly fails to unfold and reveal the structure of the three separated groups of samples.

MDS, on the other hand, preserves the sample distances in a 2D-space based on the calculation of a distance matrix (Fig. 1C,D). In ecology, distance (or dissimilarity) matrices are a major way to transpose the ecological information of samples in terms of their species composition and abundance ^40,41^. In this article we will consider classical MDS (which uses Euclidean distance and is in practice equivalent to PCA ^42,43^), and non-metric MDS (NMDS) obtained according to Sammon’s Mapping ^44^. In the latter, the elements of the multivariate space are mapped onto a lower dimensional space while retaining the original inter-point dissimilarities, by means of a nonlinear, but monotonic transformation (Sammon Mapping). Since it respects the ranking of dissimilarities, it tends to linearize the relationships between the samples. In addition, MDS will be performed also according to Bray-Curtis (MDSbc) dissimilarity and weighted UniFrac (MDSwUF) distance because they are considered the reference in metagenomics studies. Bray-Curtis dissimilarity quantifies how dissimilar two sites (samples) are based on counts (bacterial abundances), where 0 means two samples are identical and 1 means that the two samples do not share any taxa ^45,46^. Dissimilarly, the UniFrac distance, either unweighted (qualitative) or weighted (quantitative), is the most popular phylogenetic distance measure for the microbial community diversity between different samples (also known as β-diversity ^47^) and, differently from the previous discussed methods, uses the phylogenetic information (which is an external knowledge not contained in the dataset) on the taxa to compare samples. In particular, its weighted-version weights the branches of a phylogenetic tree based of the taxa abundance information ^48–51^. Hence the weighted UniFrac distance directly accounts for differences in the abundance of different kinds of bacteria, and can be crucial to describe community changes ^49^ in the studied samples.

We need to specify that both MDSwUF and NMDS are in practice nonlinear methods and weighted UniFrac is not a classical unsupervised technique like the others. In fact, MDSwUF adopts a distance that combines the information given by the bacterial abundance of the dataset with the supervised prior (external) knowledge regarding the known hierarchical phylogenetic relationship among the bacteria. However, like PCA, MDS can fail to detect patterns if data are not properly linearized ^52^. For instance, see Fig. 1C-D where MDSbc and NMDS respectively fail to resolve the Tripartite-Swiss-Roll dataset. When we consider clinical metagenomic data, this failure potentially reduces the chances of correctly pinpointing samples which may represent clinical subspecies, and thus remain undetected and undiagnosed. In brief, these methods are not efficient to perform *hierarchical embedding* directly from the abundance value, since hierarchies preserve tree-like structures, and tree-like structures follow a hyperbolic, thus nonlinear, geometry ^53–55^. Only MDSwUF is able to account for nonlinear hierarchical organization, yet this is not directly inferred from the abundance values, but rather forced as a constraint of prior supervised knowledge on the phylogeny of bacteria. For this reason we cannot offer a test on the Tripartite-Swiss-Roll dataset.

In our analysis of the Paroni Sterbini dataset, we also showed the results of a supervised technique, Linear Discriminant Analysis (LDA), which uses the labels to perform dimension reduction. LDA aims to separate the samples into groups based on hyperplanes and describe the differences between groups by a linear classification criterion that identifies decision boundaries between groups ^34^. This technique is not congruous (and sometimes statistically invalid) for small sample size datasets. The reason is that given the reduced sample size we cannot divide the dataset in a training and test set, which is a fundamental requirement of supervised methods such as LDA.

### Minimum Curvilienar Embedding (MCE)

In 2010, Cannistraci *et al.* ^23^ introduced the centred version of Minimum Curvilinear Embedding (MCE), which provided notable results in: i) visualisation and discrimination of pain patients in peripheral neuropathy, and the germ-layer characterisation of human organ tissues ^23^; ii) discrimination of microbiota in sea sponges ^36^; iii) embedding of networks in the hyperbolic space ^54^; iv) stage identification of embryonic stem cell differentiation based on genome-wide expression data ^56^. In this fourth example, MCE performance ranked first on 12 different tested approaches (evaluated on 10 diverse datasets). More recently in 2013 ^30^, the non-centred version of the algorithm, named ncMCE, has been used: i) to visualise clusters of ultra-conserved regions of DNA across eukaryotic species ^57^; ii) as a network embedding technique for predicting links in protein interaction networks ^30^, outperforming several other link prediction techniques; iii) to unsupervisedly reveal hidden patterns related with gender difference and metabolic-disease risk-factors in lipidomic profiles extracted from human plasma samples ^58^; iv) to unsupervisedly infer and visualize phylogenetic (hierarchical) relations directly from individual SNP profiles in human population genetics ^39^. Finally, also applications in non-biological problems such as the unsupervised discrimination of bad from good radar signals ^30^, represented a proof of concept of the universality of MCE for addressing nonlinear investigation of data and signals in general. Also in the case of the metagenomics studies targeting sea sponges, ^36,37^, both MCE and its non-centred variant ^23,30^ once again proved successful in detecting structure where PCA and MDS could not, or hardly find any. This is mainly because MCE/ncMCE are unsupervised and parameter-free topological machine learning for *nonlinear* dimensionality reduction and multivariate analysis, that are able to perform a *hierarchical embedding*.

This study stems from the intuition that MCE/ncMCE analysis could successfully reveal undetected patterns also in esophageal and gastric metagenomics data, where only unsupervised linear methods or classical nonlinear methods such as NMDS and MDSwUF had been used and had failed to achieve any clear-cut result ^21,22^.

Minimum Curvilinearity (MC) ^23^, the principle behind MCE and ncMCE, was invented with the aim to reveal nonlinear data structures also, and especially, in the case of datasets with few samples and many features. MC principle suggests that curvilinear (nonlinear) distances between samples may be estimated as pairwise distances over their Minimum Spanning Tree (MST), constructed according to a selected distance (Euclidean, correlation-based, etc.) in a multidimensional feature space (here the metagenomic data space). In this study, we considered Pearson-correlation based distance (refer to ^23^ for details on the way to compute the distance for the MST). The collection of all nonlinear pairwise distances forms a distance matrix called the MC-distance matrix or MC-kernel, which can be used as an input in algorithms for dimensionality reduction, clustering, classification and generally in any type of machine learning. In MCE and ncMCE, the MC-kernel (which is non-centred for ncMCE) is followed by dimensionality reduction using singular value decomposition (SVD), and then by the projection of the samples onto a two-dimensional space for visualisation and analysis. Thus, MCE/ncMCE is a form of nonlinear and parameter-free kernel PCA ^30^. In the rest of the article we will simply use the name MCE to indicate both MCE and ncMCE, since the centring transformation is related to the specific data pre-processing and will be specified for each dataset as a technical detail in the respective results’ tables.

### MCE to unsupervisedly infer and visualize phylogenetic (hierarchical) relations

A previous study by Alanis-Lobato *et al.* ^39^ showed that MCE is automatically able to unsupervisedly infer and visualize phylogenetic (hierarchical) relations directly from individual SNP profiles in human population genetics. Precisely, ncMCE detected separation between ethnic groups and provided an ordering over the discriminating dimension that was related to the phylogenetic organization of these populations.

This ability of MCE to infer and visualize phylogenetic (hierarchical) relationships was confirmed in our study on the Paroni Sterbini *et al.* dataset ^22^ (see Results section-‘ *Gastric tissue dataset unsupervised analysis’*). As previously mentioned (see the previous section ‘*PCA, MDS (or PCoA) and LDA’*), MDSwUF uses a weighted Unifrac distance that combines the prior knowledge of the bacterial phylogenetic tree with the information given by the bacterial abundance. Here we show that MCE perform better than MDSwUF on the Paroni Sterbini *et al.* dataset, due to its ability to infer the (hierarchical) phylogenetic relationship among the bacteria directly from the bacterial abundance of the dataset, by performing a hierarchical embedding. Hence, MCE can be used to compare the composition of microbial communities in the studied samples, where the phylogenetic information is instead directly inferred from bacterial abundance, differently from MDSwUF.

### Procedure to evaluate the performance of the dimension reduction algorithms

The performance of the mentioned dimension reduction algorithms is evaluated as the ability to separate the samples in the first two dimensions of embedding since, as discussed above, this is one of the preferred unsupervised strategies to investigate the presence of patterns in multidimensional datasets. In order to quantitatively evaluate the performance, we use a recently proposed index for sample separation ^59^. This index can be defined for any separation-measure and in this study we considered three well-known measures: p-value of Mann-Whitney U test, Area Under the ROC-Curve (AUC) and Area Under the Precision-Recall curve (AUPR), that are regularly used to quantitatively measure the performance of a binary predictor.

More precisely, in the 2D space a line is drawn between the centroids of the two groups that are compared, subsequently all the points are projected on this line and then a p-value, AUC and AUPR are computed for the projected points. This new index is named *projection-based separability index* (PSI) and can actually be applied not only in a 2D space, but in any N dimensional space. For the calculation of the centroids we consider the 2D-median of each cluster/class’s group. In case more than two groups are present in a dataset, all the p-values, AUC and AUPR between the possible pair-groups are computed, and the average values of all the pairwise p-values, AUC and AUPR are chosen as an overall estimator of separation between the groups in the 2D reduced space. This case applies only to the Paroni Sterbini dataset, which is composed of three or, possibly, four groups of samples. All the other datasets are instead composed of two groups.

It is important to note that the PSI was also applied to the data in the original high-dimensional (HD) space, as a reference to see how good the unsupervised dimension reduction approaches are in preserving the original group separability of the HD space.

All the algorithms were tested considering (when allowed by the dimension reduction method) data centring or non-centring. In addition, multiple normalization options were investigated and the datasets were considered under a certain type of normalization: division by the column - which reports the OTU - sum (indicated by DCS); division by the row – which reports the sample - sum (indicated by DRS); function log10(1+x) applied to the dataset (indicated by LOG).

### From Markov Clustering (MCL) to Minimum Curvilinear Markov Clustering (MC-MCL)

MCL is an unsupervised algorithm for the clustering of weighted graphs based on simulations of (stochastic) flow in graphs ^60^ (http://micans.org/mcl/). By varying a single parameter called inflation (with values between 1.1 and 10), clustering patterns on different scales of granularity can be detected. For clustering samples of a multidimensional dataset, the workflow starts with the computation of correlations (generally Pearson correlations) between the samples, and creates an edge between each pair of samples, where the edge-weight assumes the value of the respective pairwise positive sample correlation, or values zeros in case of negative correlations. This generates a weighted correlation graph (network), which is used as a map to simulate stochastic flows and detect the structural organization of clusters in the graph.

With the purpose of creating and testing a nonlinear variant of the MCL algorithm, we adopt an innovative algorithm which was recently proposed and called MC-MCL ^61^. The idea is the following. The MC-kernel – discussed above in the MCE section - is a nonlinear distance matrix (or kernel) that expresses the pairwise relations between samples as a value of distance: small samples distance indicates sample similarity, while large samples distance indicates sample dissimilarity. Here we reverse (using the following function: *f*(*x*) = 1 − *x*) and after this we put to zero the negative values of the *MC-distance* kernel to get a *MC-similarity* kernel, where small values (close to zero) indicate low sample similarity and large values (close to one) indicate high sample similarity. A technical detail: for the computation of the MC-distance kernel, it is necessary to firstly square root the original distances (correlation-based) between the samples. As already investigated in ^23^, this attenuates the estimation of large distances and amplifies the estimation of short distances; consequently it helps to regularize the nonlinear distances inferred over the MST in order to subsequently use them for message passing ^23^ (such as affinity propagation) or flow simulation (such as MCL) clustering algorithms.

Then, the standard stochastic flow simulations of MCL algorithm runs on the graph weighted with the values of the MC-similarity kernel (which collects pairwise *nonlinear* associations between samples) instead of the Pearson-correlation kernel (which collects pairwise *linear* associations between samples). In practice, this is a new algorithm for clustering that is a nonlinear version (based on the MC-kernel) of the classical MCL. The goal of the MC-MCL analysis is to verify whether the use of the MC-kernel improves performance, by solving nonlinearity, not only in dimension reduction (such as in MCE) but also in clustering (such as in MC-MCL).

### Procedure to evaluate the performance of clustering algorithms

The clustering algorithms MCL and MC-MCL were applied to the datasets, either raw, or after the same normalization procedures used before dimensionality reduction (DCS: division by column (OTU) sum; DRS: division by row (sample) sum; LOG: function log10(1+x) applied to the dataset) and their performance was evaluated by means of accuracy. The accuracy is computed as the ratio of the number of samples assigned to the correct clusters over the total number of samples. For both MCL and MC-MCL, we tested Pearson and Spearman correlations to build the similarity measure to feed into the clustering methods. The Spearman correlation can also detect a subclass of nonlinear associations (which have monotonic shape function) or correct for outliers. Differently from what suggested for large gene datasets with thousands of samples in ^60^ (http://micans.org/mcl/), in this study we had to consider the whole set of original positive correlations without applying any threshold (cut-off) to the values. This was compulsory, since we considered datasets with few samples. In our case, to keep the graph connected, with one unique connected component, we could not introduce any kind of threshold that would otherwise alter the real graph connectivity (dividing the graph in disconnected components) and hence the clustering result. Since the MCL algorithm needs a single input parameter (inflation) to control the granularity of the output clustering, we ran it for different inflation values until we achieved the desired number of clusters. Finally, in the Paroni Sterbini *et al.* dataset ^22^ it was not clear in advance whether the correct number of clusters present in the multidimensional space was three or four. Hence, we tested the clustering algorithms considering as output both three and four clusters’ configurations, and we identified as the best solution the one that offered the highest accuracy.

### PC-corr network

Furthermore, we investigated the effect of PPI on the microbiota of gastric fluid and gastric mucosa in dyspeptic patients, and the changes induced by *H. pylori* infection on the gastric mucosal microbiota, by means of the PC-corr approach ^62^. PC-corr represents a simple algorithm that associates to any PCA segregation a discriminative network of features’ interactions ^62^. It is a method for linear multivariate-discriminative correlation network reverse engineering, that, thanks to its multivariate nature, can help to stress and squeeze out the underlying combinatorial and multifactorial mechanisms that generate the differences between the studied conditions ^62^. Hence, for the studied datasets, it can be employed to point out the possible presence of bacterial alterations and their interplay, induced by a medical treatment (PPIs in dyspepsia) or infectious state (*H. pylori*).

### Computing platforms adopted to implement the algorithms

Dimensionality reduction was performed in MATLAB on the abundance matrix of genus-level taxonomic assignments, with samples in rows and taxonomic assignments (OTUs) in columns. For MDSwUF, the computation of the weighted UniFrac distance was performed in R. We used the following MATLAB functions to calculate PCA, MDS and NMDS (Sammon Mapping) respectively: *svd*, *cmdscale* and *mdscale*. For the calculation of Bray-Curtis dissimilarity, we used the function MATLAB *f_braycurtis* in the Fathom Toolbox ^63^ (http://www.marine.usf.edu/user/djones/matlab/matlab.html). Instead, for the calculation of the weighted Unifrac distance for all sample pairs, we used the R function *UniFrac* in the phyloseq package (https://bioconductor.org/packages/release/bioc/html/phyloseq.html), after creating a phyloseq-class object (with R function *phyloseq* in the same package) that contains both the abundance table (OTU table) and the phylogenetic tree. The MATLAB code for MCE/ncMCE is available online at: https://sites.google.com/site/carlovittoriocannistraci/5-datasets-and-matlab-code/minimum-curvilinearity-ii-april-2012. For MCL clustering, we installed the MCL-edge software (http://micans.org/mcl/) in a Windows environment, following the procedure suggested by the authors in the software website. To apply this algorithm, we created a MATLAB function that generates automatically the input for MCL (equivalent to the mcxarray function in the software) and then uses a system call to run MCL in a UNIX-like environment (Cygwin, https://www.cygwin.com/). PC-corr method was performed in MATLAB on the abundance matrix of the genus-level taxonomic assignments, with samples in rows and taxonomic assignments in columns. The PC-corr algorithm is available as MATLAB function (as well as R function) at: https://github.com/biomedical-cybernetics/PC-corr_net. Then the obtained PC-corr networks were displayed by Cytoscape (http://www.cytoscape.org/).

## Results

To answer the five questions stated in the Background section, we analysed the abovementioned 16S rRNA gene sequencing datasets with information on PPI consumption in dyspeptic patients, following the workflow shown in Fig. 2. It is important to underline that, in one of the three initially analysed datasets (in Paroni Sterbini *et al.*^22^), we have the additional information on positivity or negativity to *H. pylori* infection. A fourth dataset (Parsons *et al.* ^29^) is used only for the validation of the PC-corr network results and it contains not only information on PPI consumption but also additional information on positivity or negativity to *H. pylori* infection.

**Figure 2.**
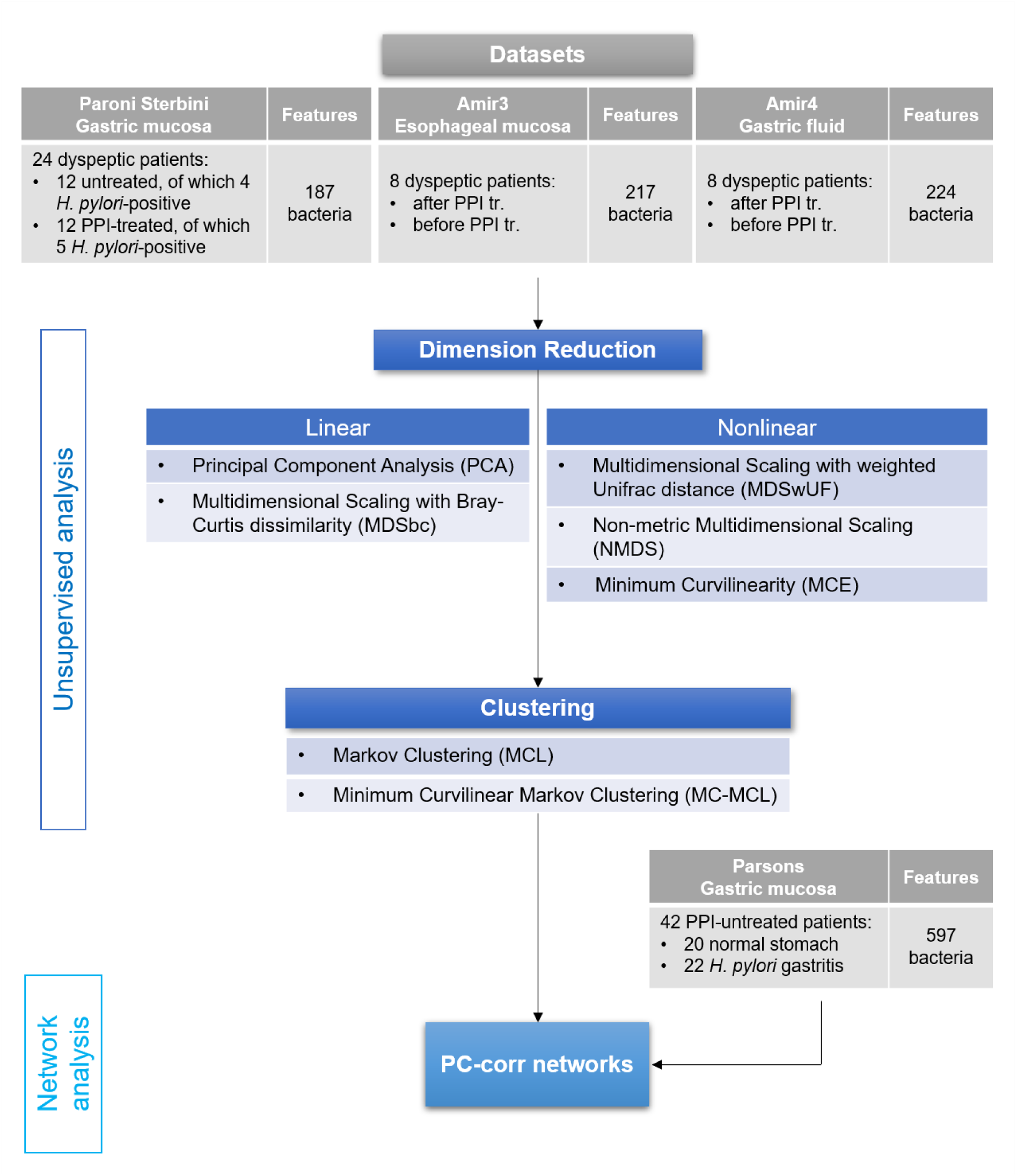
Flowchart of the data analysis. To answer the five questions under investigation in our study, we implemented a workflow based on machine learning tools. Following the flowchart shown in the figure, we analysed three 16S rRNA gene sequencing datasets with information on PPI use in dyspeptic patients; for one of the datasets (Paroni Sterbini *et al.* ^22^), patients were also determined to be positive or negative to *H. pylori* infection. Firstly, we performed unsupervised dimension reduction, both linear and nonlinear, in the first two dimensions of embedding. Nonlinear dimension reduction will show the presence of hidden patterns, in the form of sample groups. Secondly, nonlinear clustering was applied to confirm the well-possedeness of the hidden patterns found by nonlinear dimension reduction. Lastly, our workflow ends with the PC-corr algorithm, that reveals which combination of bacteria (features) are responsible for the identified differences between the groups of samples. A fourth dataset (Parsons *et al* ^29^.) is used only for the validation of the PC-corr network results and it contains information of PPI treatment and *H. pylori* infection.

Unsupervised approaches were chosen for dimension reduction, and clustering because supervised (constrained) methods have been shown to perform poorly on small datasets, as explained in the paper by Smialowski *et al.* ^31^ and the work by Zagar and colleagues ^56^.

Firstly, we performed unsupervised dimension reduction, both linear and nonlinear (described in the *‘Methods-PCA, MDS (or PCoA) and LDA’ and ‘Methods-Minimum Curvilinear Embedding (MCE)’*) and we focused on the first two dimensions of embedding as it is common practice ^25^. As we will show, linear techniques will fail to bring out the patterns in the microbial datasets, related to PPI-treatment. Instead, nonlinear dimension reduction will reveal the presence of hidden patterns related to PPI treatment. In particular, in the gastric biopsies dataset (Paroni Sterbini *et al.* ^22^), nonlinear dimension reduction will point out the evidence of PPI perturbation. Secondly, clustering algorithms were applied to the studied datasets to confirm that the hidden patterns detected by nonlinear dimension reduction are well posed. Finally, the PC-corr algorithm ^62^ is used to find the bacteria community (features) that make the difference between the patterns or groups, allowing our understanding of the PPI-induced and *H. pylori*-induced microbial perturbations.

### Gastric tissue dataset unsupervised analysis

According to the questions formulated in our study, we are interested in an unsupervised approach to verify whether PPI drugs cause a major change in the gastric tissue microbiota of dyspeptic patients regardless of the initial pathological infection due to *H. pylori* ^22^.

In our first analysis, we focused on the Paroni Sterbini *et al.* dataset ^22^ and, to facilitate the visualization of the sample separations in the 2D reduced space, we assigned: red colour to untreated dyspeptic patients without *H. pylori* infection (HP-); green colour to untreated dyspeptic patients with *H. pylori* infection (HP+); and blue colour to patients treated with PPI regardless of their *H. pylori* infection (PPI). However, to help to detect also the effect of the *H. pylori* infection we reported the labels close to each sample, with a ‘+’ indicating the infection (PPI+) or a ‘-’ indicating the absence of infection (PPI-). Finally, we also tested whether this separation into three main groups (HP-, HP+, PPI) is more truthful, from the metagenomics data standpoint, than the one in four groups (HP-, HP+, PPI-, PPI+).

Figure 3 shows the results of the multivariate techniques widely employed in metagenomic studies, PCA (Fig. 3A), MDSbc (Fig. 3B) and MDSwUF (Fig. 3C), and NMDS (with Sammon Mapping) (Fig. 3D) (for more detail see the corresponding method section; the plots represents the best results based on average p-value in Supplementary Table S1), which could only differentiate the group of untreated *H. pylori* positive samples (green dots) with respect to the group of untreated *H. pylori* negative samples (red dots), and no further separation is significantly detectable. Considering the PSI results, the p-values are significant (p-value<0.05, Table 1 and Fig. 3) (evaluated in the 2D embedding space, for details see ‘*Procedure to evaluate the performance of the dimension reduction algorithms’*). PCA and NMDS exhibit the lowest p-value (0.0090), while MDSwUF and MDSbc displays p-values higher than 0.01 (respectively 0.011 and 0.021). This trend is also confirmed by their AUC and AUPR values, with highest values for PCA (AUC=0.924, AUPR=0.960) and NMDS (AUC=0.924, AUPR=0.954). Indeed, in all the plots there is a visible trend of separation between PPI-treated (blue dots) and untreated (red and green dots) samples, but this is not sufficient to declare the presence of the complete separation, and a manifest ‘crowding problem’ ^30^ mixes the two cohorts together. According to this output, the dataset appears to be strongly influenced by the presence of *H. pylori*, which is the predominant taxon (abundance > 50%, Supplementary Table S2, percent abundance sheet) in four of the untreated *H. pylori* positive patients: where *H. pylori* is predominant, sample groups are quite close to one another and far from all the other samples in all four multivariate analyses (Fig. 3). Thus, PCA and MDS mainly show us that these metagenomes separate according to *H. pylori* abundance, and there is no treatment-related pattern.

**Figure 3.**
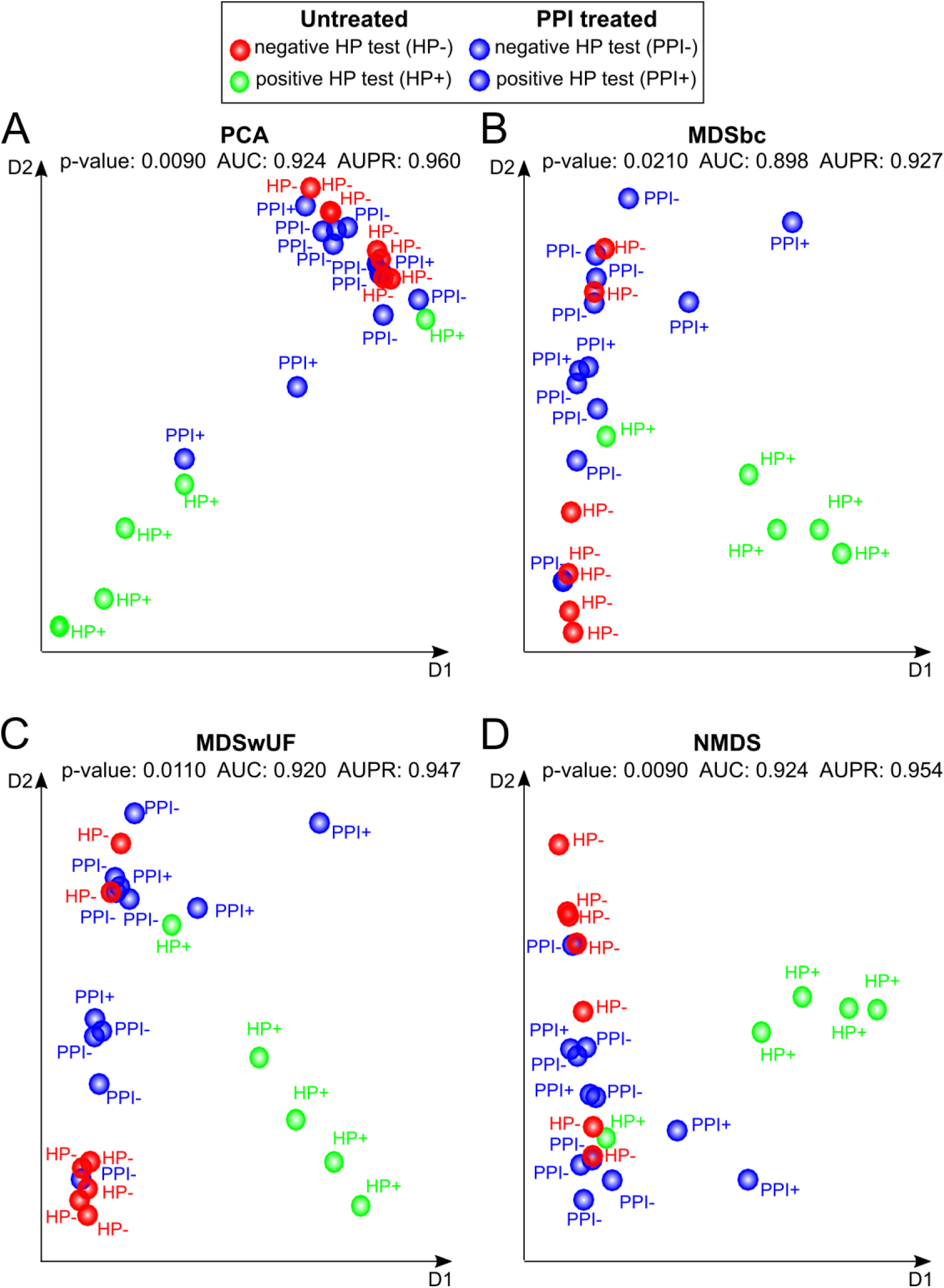
Dimension reduction techniques usually employed in metagenomic data analysis and applied to the Paroni Sterbini dataset. The plots represent the best PCA and MDS results based on (average) p-value projection-based separability index (PSI) for the three different labels (PPI-treated, untreated HP+ and untreated HP-), evaluated in the 2D embedding space. Moreover, also the average values of all pairwise AUC and AUPR PSI are reported as overall estimators of separation between the groups in the 2D reduced space. A) PCA; B) MDS with Bray-Curtis dissimilarity (MDSbc); C) MDS with weighted UniFrac distance (MDSwUF); D) non-metric MDS with Sammon Mapping (NMDS). Blue dots represent PPI-treated samples, while red and green dots are the untreated samples which resulted either negative (red) or positive (green) to the *H. pylori* test (histological observation and urease test).

**Table 1.**
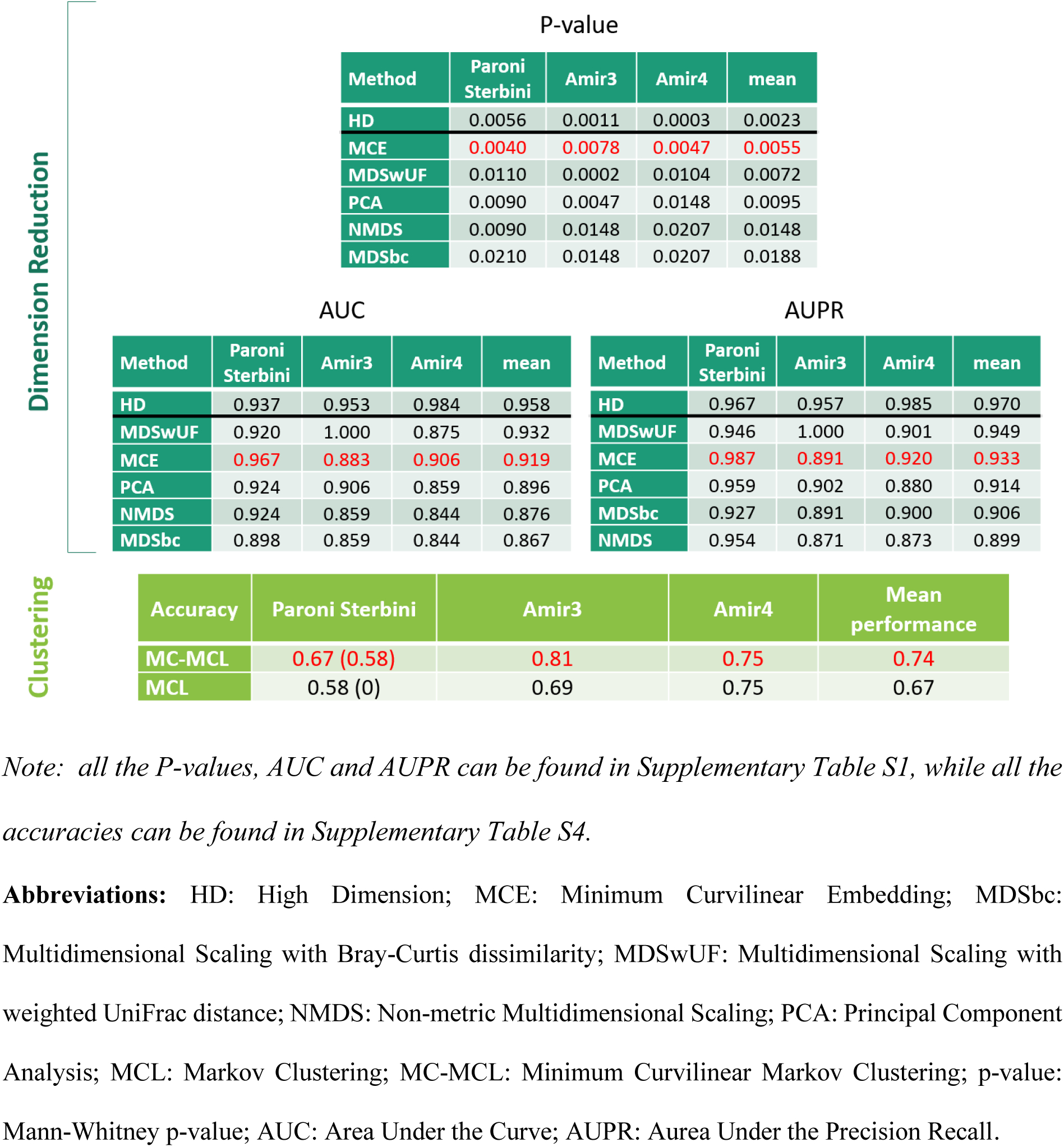
Results of unsupervised analysis on the original datasets. Best results of unsupervised dimension reduction techniques (top panel) and of clustering (bottom panel). (**Top panel**): Best results of unsupervised dimension reduction techniques according to the index for sample separation in the space of the first two dimensions of embedding. HD (no dimension reduction) represents the reference results to see how good the separability present in the high dimensional space is preserved by dimension reduction techniques. Results are ordered from the best (top) to the worst (bottom) method. For the Paroni Sterbini dataset, we show the results for three different labels (PPI-treated, untreated HP+ and untreated HP-). For the Amir datasets, the p-values were computed for two groups, identified by the presence or absence of PPI treatment. (**Bottom panel**): Best results of clustering (highest accuracies, regardless of the normalization and type of correlation) MCL and MC-MCL, in each of the three studied datasets (Paroni Sterbini, Amir3 and Amir4), and the mean performance (mean of the highest accuracies) across all the datasets. For Paroni Sterbini dataset, we show the results for three clusters (PPI-treated, untreated HP+ and untreated HP-) and in brackets the results for four clusters (PPI-treated HP+, PPI-treated HP-, untreated HP+ and untreated HP-). Instead, for Amir datasets, the accuracies were computed for two groups, identified according to the presence or absence of PPI treatment.

Non-centred MCE (Figure 4A, DCS normalization) was the best performing technique, with a p-value of 0.004, AUC of 0.967 and AUPR of 0.987 (Table 1) (for details see Supplementary Table S1). It even outperforms the nonlinear methods NMDS (Sammon Mapping) and MDSwUF, since it is automatically able to infer the (hierarchical) phylogenetic relationship among the bacteria directly from the bacterial abundance of the dataset by performing a hierarchical embedding, as already shown in the study of Alanis-Lobato *et al.* ^39^ (see *‘Methods-MCE to unsupervisedly infer and visualize phylogenetic (hierarchical) relations*’). Furthermore, the MCE performance does not depend on its centring/non-centring, in fact the centred MCE version resolves the nonlinearity in the data too. Whereas, PCA regardless of being centred or non-centred does not resolve the nonlinearity in the data.

**Figure 4.**
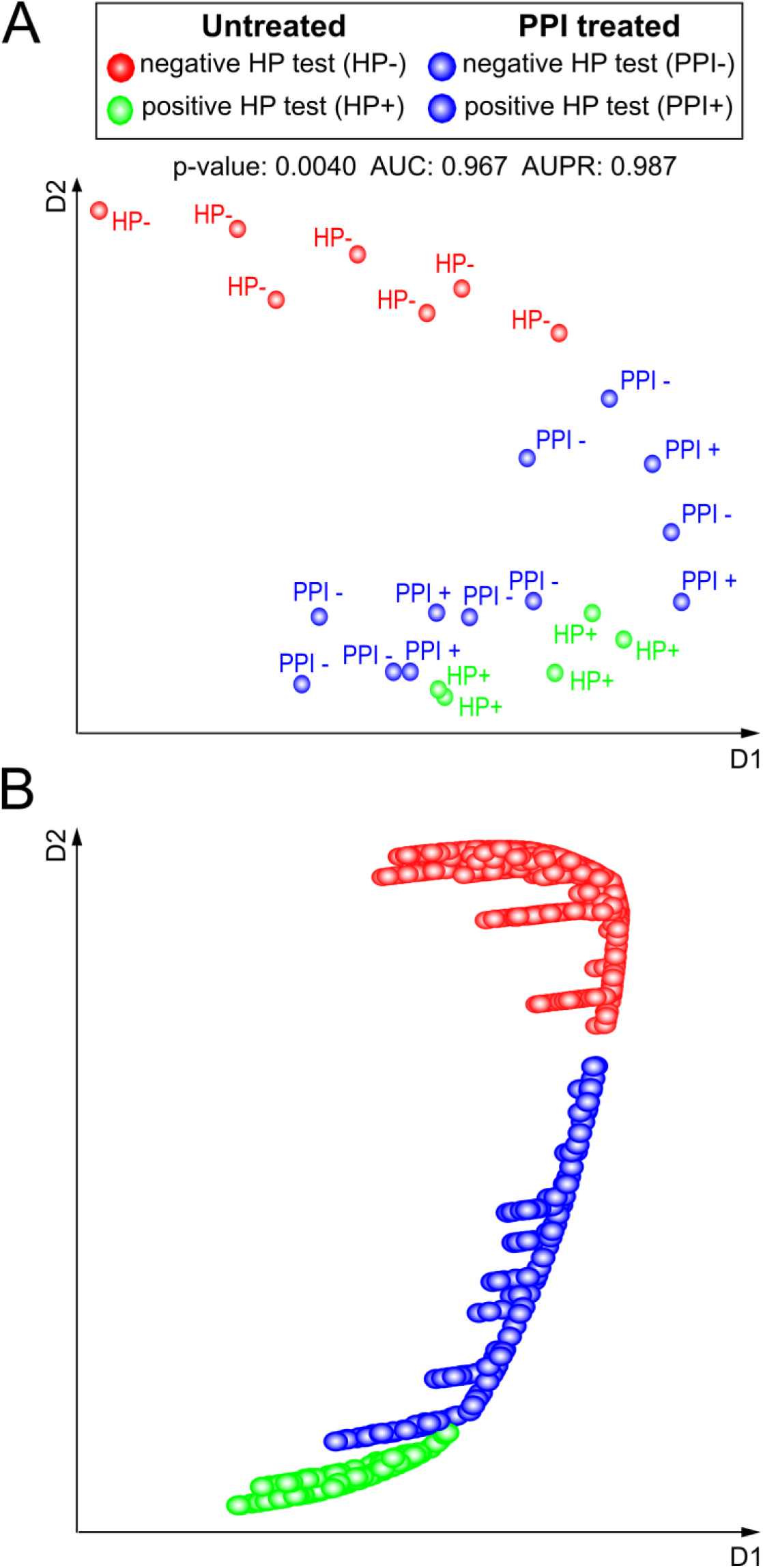
MCE, a topological machine learning for nonlinear and hierarchical dimension reduction. (**A**) Results on the Paroni Sterbini *et al.*^22^ dataset. The shown best MCE result is based on (average) p-value projection-based separability index (PSI) for the three different labels (PPI-treated, untreated HP+ and untreated HP-), evaluated in the 2D embedding space under the DCS normalization. The average values of all pairwise AUC and AUPR PSI are reported as well as overall estimators of separation between the groups in the 2D reduced space. Blue dots represent PPI-treated samples, while red and green dots are the untreated samples which resulted either negative (red) or positive (green) to the *H. pylori* test (histological observation and urease test). **(B**) Results on the Tripartite-Swiss Roll. The three different colours (red, blue and green) represent the three partitions of the Swiss-roll manifold.

While MDS and PCA are confounded by the mixture of factors characterizing the samples and do not manage to resolve the differences between treated and untreated samples, non-centred MCE is the only technique that visibly separates samples by ordering them along the second dimension into three groups, detecting a treatment-related structure in the data (Fig. 4A). This is plausible, because in any non-centred embedding the first dimension points towards the centre of the manifold ^30^, while the second dimension in the case of non-centred MCE represents the direction of higher topological nonlinear extension of the manifold. Interestingly, untreated *H. pylori* negative samples (red dots, HP-) gather in the upper tail of the samples’ distribution, while treated samples (blue dots, PPI), both *H. pylori* test positive (PPI+) and negative (PPI-), are mixed and show no other internal discernible groups. Untreated *H. pylori* positive samples (green samples, HP+) gather at the bottom of the plot (Fig. 4A). Unlike the other approaches, non-centred MCE detects a treatment-related structure in the data and separates patients into three, not four, groups: PPI-treated, untreated *H. pylori* negative and untreated *H. pylori* positive. This last group appears as a subgroup marginally discriminating from the PPI-treated group and the topology of the samples seems to suggest that PPI treatment modifies the gastric microbiota of *H. pylori*-negative patients with dyspeptic symptoms and gastric mucosa inflammation, shifting their gastric ecosystem in the same direction of PPI-treated *H. pylori*-positive patients. We speculate that the fact that PPI treatment and *H. pylori* infection determine the samples to gather in a similar position (i.e. out of the PPI-untreated/HP-negative group) in the non-centred MCE reduced space, indicates that both the PPI drugs and *H. pylori* induce an ecological change in the stomach, which might be driven by similar mechanisms. As a matter of fact, *H. pylori* can colonize the acidic lumen of the stomach thanks to its ability to hydrolyse urea into carbon dioxide (CO_2_) and ammonia (NH_3_) ^64^, thus increasing the intragastric pH. On the other hand, PPIs obtain the same result through the inhibition of acid secretion in gastric parietal cells, which blocks H^+^/K^+^ -ATPases. Both processes are therefore shifting the gastric environment towards an alkaline condition. Thus, MCE provides an ordering of the groups along the second dimension that is related to pH increment (from HP- to PPI+).

Similarly to the Paroni Sterbini *et al.* microbial dataset, the Tripartite-Swiss-roll dataset (that is a synthetic dataset containing nonlinear structures obtained by tri-partitioning a discrete Swiss-Roll manifold ^38^ in a three-dimensional space, for more details see the method section: The Tripartite-Swiss-Roll dataset’), presents a hierarchical-organized nonlinearity (Fig. 1A). And also in this case, similarly to the result of the Paroni Sterbini *et al.* analysis, non-centred MCE is able to perform a hierarchical embedding that orders the hidden subgroups of the dataset along the second dimension of embedding (Fig. 4B). On the contrary - as already commented in the method section - PCA, MDSbc and NMDS (Fig. 1B-D) were unable to resolve the nonlinearity of the Tripartite-Swiss-Roll: its three partitions are either superimposed (Fig. 1B, D) or twisted in a horseshoe shape (Fig. 1C). Indeed, the Tripartite-Swiss-Roll is purposely created to reproduce a manifold that is nonlinear and discontinuous (broken in three parts) such as the results of MCE analysis of Paroni Sterbini *et al.* seems to be.

For the Paroni Sterbini dataset, we also performed a supervised linear approach for dimension reduction, LDA (Supplementary Figure S1), yet the cross-validation test showed that this constrained technique could re-assign samples to their groups with 54% of error (ldaCVErr in Supplementary Table S3), confirming its statistical invalidity for the small size dataset problem.

Moreover, the clustering algorithms MCL and MC-MCL, that is the minimum curvilinear version of MCL were applied to the Paroni Sterbini *et al.* dataset and the best results (highest accuracies) are shown in Table 1 (bottom panel) (for more details see the methods’ sections ‘*From Markov Clustering (MCL) to Minimum Curvilinear Markov Clustering (MC-MCL)’ and ‘Procedure to evaluate the performance of clustering algorithms’*). MC-MCL performs better than the MCL (both for three and four clusters), even if their accuracies are not remarkably high, confirming that difficulties in pattern-recognition arise also from the presence of three clusters in the high-dimensional space. In addition, the hypothesis of three clusters seems more congruous than four clusters, because both MC-MCL and MCL decrease their accuracies in detecting four clusters.

While MC-MCL represents the minimum curvilinear version of MCL, MCE is the minimum curvilinear version of PCA, particularly valuable for small sample size datasets. The principle behind them is MC^23^, that suggests that curvilinear (nonlinear) distances between samples may be estimated as pairwise distances over their Minimum Spanning Tree (MST) (constructed according to a selected distance). In fact, as explained in ^65^, to approximate nonlinear (curvilinear) distances between the points of the manifold it is not necessary to reconstruct the nearest-neighbour graph. Indeed, a greedy routing process (that exploits a norm, for instance Euclidean) between the points in the multidimensional space is enough to efficiently navigate the hidden network that approximates the manifold in the multidimensional space. And a preferable greedy routing strategy, at the basis of MC-kernel, is the minimum spanning tree (MST).

Overall, we can conclude that both MCE in dimensionality reduction and MC-MCL in clustering perform better than the respective non-MC-based versions, and this result confirms the presence of nonlinear complexity in this dataset, generated by a three-body interaction (presence of three clusters). In addition, when considering correlation-based distances, they do not react to the presence of compositionality, since pairwise correlations are computed between samples. Compositionality instead is a problem that arises when the correlations is computed between OTUs (features) from metagenomics abundance data (which are normalized by diving each OTU count to the total sum of counts in the sample ^66,67^), which yields unreliable results due to dependency of microbial relative abundances.

Moreover, because of the discovered major nonlinear complexity in the Paroni Sterbini gastric biopsy dataset, we wanted to verify whether it was generated by multi-grouping (three-body interaction problem associated to the presence of three hidden clusters). To do so, we applied PCA to three subsampled versions of the dataset (with the best normalization originally found for the complete dataset), each corresponding to the combination of two groups (Fig. 5A-C), and PCA could find significant separation (p-values <0.02 and AUC, AUPR > 0.80). To further confirm that the presence of multiple sample groups generates the data complexity, we did the same for the Tripartite Swiss-Roll (Fig. 5D-F), where we recovered the discrimination, even though two comparisons overlap to some extent (Fig. 5D and F). Furthermore, to have another confirmation that the PPI-treated samples are not separable for *H. pylori* infection, we analysed the dataset considering exclusively the PPI-treated samples. The result is that no internal separation related to *H. pylori* infection emerges within the PPI-treated patients, as shown by the best MCE result (Supplementary Figure S2).

**Figure 5.**
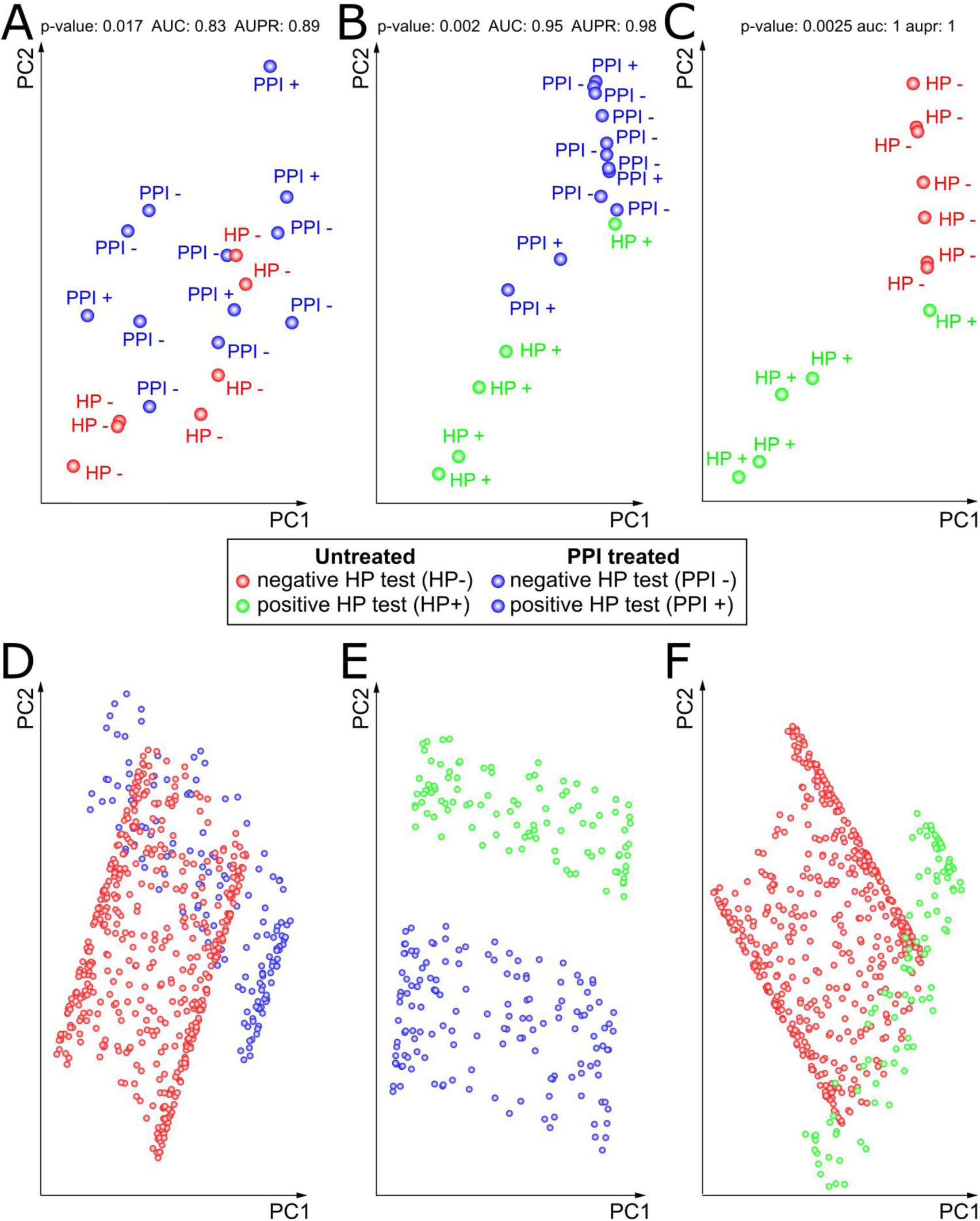
Pairwise PCA of Paroni Sterbini’s gastric samples and of the Tripartite-Swiss-Roll. **(A-C)** PCA was applied to three subsampled versions of the Paroni Sterbini dataset (keeping the best normalization found for the original dataset), each corresponding to the combination of two groups: A) PPI-treated and untreated *H. pylori* negative samples; B) PPI-treated and untreated *H. pylori* positive samples; C) untreated *H. pylori* negative and untreated *H. pylori* positive samples. The p-value, AUC and AUPR PSI are reported as well as overall estimators of separation between the groups in the 2D reduced space **(D-F)** In a similar manner, PCA was applied to the three datasets obtained by subsetting the Swiss-roll dataset, each one corresponding to a combination of two groups: D) red vs blue groups, E) blue vs green groups, F) red vs green groups.

In conclusion, the results confirm that linear techniques, even if supervised like LDA, are not able to resolve the differences in the data due to the presence of nonlinear complexity generated by the three-body interaction (HP-, HP+ and PPI). Once the complexity is reduced to a two-body interaction, the problem tends to vanish and PCA can detect significant differences between the groups, as shown by the PCA pairwise comparisons.

Hence, the results of unsupervised analysis on Paroni Sterbini *et al.* dataset show that PPI treatment causes a major change in gastric mucosal communities of dyspeptic patients, regardless of the initial pathological infection due to *H. pylori*.

### Comparison of unsupervised analysis in three gastro-esophageal datasets

We compared the performance of unsupervised analysis (dimensional reduction and clustering) in the Paroni Sterbini dataset ^22^ (gastric biopsies) and two additional datasets by Amir and colleagues ^21^, that investigated the PPI influence on the esophageal microbiota (Amir3) and gastric fluid (Amir4).

Table 1, top panel, shows the best results in performance of unsupervised dimension reduction (PCA, MDSwUF, MDSbc, NMDS, MCE, for details see *‘Methods - PCA, MDS (or PCoA) and LDA’ and ‘Methods - Minimum Curvilienar Embedding (MCE)’*) according to the PSI (projection-based separability index) in the space of the first two dimensions of embedding, based on the p-value of Mann-Whitney U test, AUC and the AUPR, on the three different datasets (for more details on the PSI see *‘Methods - Procedure to evaluate the performance of the dimension reduction algorithms’*). The mean performance across all datasets is shown in the last column of the table for each method. The corresponding ranked performance for each method, based on p-value, AUC and AUPR, is presented instead in Table 2. For the Paroni Sterbini dataset, we show the results for three different labels (untreated HP-, untreated HP+ and PPI-treated). For the Amir datasets, the p-values were computed for two groups, identified by the presence or absence of PPI treatment. The PSI was also applied to the data in the original high-dimensional (HD) space, as a reference to see how good the unsupervised dimension reduction approaches are in preserving the group separability in the HD. Moreover, the average p-value, AUC and AUPR best results with standard error on the original datasets, when applying leave-one-out-cross-validation (LOOCV), are shown in Supplementary Table S5.

**Table 2.** Ranked performance of unsupervised dimension reduction techniques on the original datasets. The table shows the ranked performance of unsupervised dimension reduction techniques according to the index for sample separation (based on Mann-Whitney P-value, AUC and AUPR) in the space of the first two dimensions of embedding, for the three studied datasets (Paroni Sterbini, Amir3 and Amir4). Each rank is related to the results obtained in Table 1, top panel. The results are ordered by the mean performance (fourth column) from the best (top) to the worst (bottom) method.

| P-value |  |  |  |  |
| --- | --- | --- | --- | --- |
| Method | Paroni Sterbini | Amir3 | Amir4 | mean |
| HD | 2 | 2 | 1 | 1.67 |
| MCE | 1 | 4 | 2 | 2.33 |
| MDSwUF | 4 | 1 | 3 | 2.67 |
| PCA | 3 | 3 | 4 | 3.33 |
| NMDS | 3 | 5 | 5 | 4.33 |
| MDSbc | 5 | 5 | 5 | 5 |

| AUC |  |  |  |  | AUPR |  |  |  |  |
| --- | --- | --- | --- | --- | --- | --- | --- | --- | --- |
| Method | Paroni Sterbini | Amir3 | Amir4 | mean | Method | Paroni Sterbini | Amir3 | Amir4 | mean |
| HD | 2 | 2 | 1 | 1.67 | HD | 2 | 2 | 1 | 1.67 |
| MCE | 1 | 4 | 2 | 2.33 | MCE | 1 | 4 | 2 | 2.33 |
| MDSwUF | 4 | 1 | 3 | 2.67 | MDSwUF | 5 | 1 | 3 | 3 |
| PCA | 3 | 3 | 4 | 3.33 | PCA | 3 | 3 | 5 | 3.67 |
| NMDS | 3 | 5 | 5 | 4.33 | MDSbc | 6 | 4 | 4 | 4.67 |
| MDSbc | 5 | 5 | 5 | 5 | NMDS | 4 | 5 | 6 | 5 |

**Abbreviations:** HD: High Dimension; MCE: Minimum Curvilinear Embedding; MDSbc:
Multidimensional Scaling with Bray-Curtis dissimilarity; MDSwUF: Multidimensional Scaling with
weighted UniFrac distance; NMDS: Non-metric Multidimensional Scaling; PCA: Principal Component
Analysis; p-value: Mann-Whitney p-value; AUC: Area Under the Curve; AUPR: Aarea Under the
Precision Recall.

For the Paroni Sterbini dataset, the PSI evaluation in the first two dimensions of embedding identifies MCE as the best dimension reduction technique that is able to preserve the group separability in the HD space. Surprisingly, MCE (presented in Fig. 4A, p-value= 0.0040, AUC = 0.967, AUPR=0.987) outdoes HD in sample separation in three groups (for HD, p-value= 0.0056, AUC= 0.937, AUPR=0.967). Similarly, in Amir4, MCE (p-value=0.0047, AUC=0.906, AUPR=0.920) succeeds in preserving the separability of the original HD space (in HD, p-value=0.0003, AUC=0.984, AUPR=0.985), better than the other dimension reduction methods. Finally, dimension reduction analysis on the Amir3 dataset shows that esophageal biopsies were significantly different before and after PPI treatment, as shown by MDSwUF results (p-value= 0.0002, AUC=1=AUPR), that surpass the p-value, AUC and AUPR values in HD space (p-value=0.0011, AUC=0.953, AUPR=0.957). Markedly, MDSwUF reaches a value of AUPR and AUC of 1, meaning perfect classification of the samples.

Overall, when averaging across all datasets, the two metrics based on AUC and AUPR pointed out that MDSwUF (AUC=0.932, AUPR= 0.949) gave the best results of separability compared to HD (AUC=0.958, AUPR=0.970), followed by MCE with closer results (AUC=0.919, AUPR=0.933), while MCE gave the highest separability according to p-value (p-value=0.0055). Then PCA is the third best result (p-value=0.0095, AUC=0.896, AUPR=0.914), followed by NMDS and MDSbc. However, to conclude what is the best method, we considered an evaluation based on ranking (Table 2). It is important to note that MCE was the dimension reduction approach that ranked first in performance across all the datasets, followed by MDSwUF (Table 2). Hence, the results of sample separability suggest the presence of hidden patterns that emerge by applying nonlinear dimension reduction techniques like MCE and MDSwUF.

Then clustering algorithms, MCL and its Minimum Curvilinear version (for more information see *‘Methods - From Markov Clustering (MCL) to Minimum Curvilinear Markov Clustering (MC-MCL)’*), were used to confirm the well-possedeness of the hidden patterns that were recognized by nonlinear dimension reduction. The best results as highest accuracies in each dataset and the mean performance across all the datasets are exhibited in Table 1, bottom panel. As already discussed in the previous section, the minimum curvilinear version of MCL (MC-MCL, acc=0.67) outperforms the MCL clustering algorithm (acc=0.58) in the Paroni Sterbini dataset, confirming the presence of underlying non-linear complexity in the data. However, the accuracy doesn’t reach high values, because of the difficulty in pattern recognition generated by the three-body problem in the HD space. Curiously, the accuracies for four clusters (HP-, HP+, PPI-, PPI+) drop to 0.58 for MC-MCL and to 0 for MCL, supporting the hypothesis that three clusters are more congruous than four clusters. Notably in Amir3, MC-MCL attains high clustering accuracy (acc=0.81), compared to MCL (acc=0.69). This is the dataset for which, surprisingly, Amir and collaborators did not find significant changes in the esophageal tissue microbiota following PPI-treatment, using classical MDS unsupervised multivariate method with unweighted UniFrac distance ^21^. Instead, in the gastric fluid dataset (Amir 4), MC-MCL and MCL got the same accuracy of 0.75, where a significant separation of samples according to PPI consumption was already proved in the original article ^21^.

However, we have to clarify that normalizations besides scaling (DRS and DCS) and log-transformation (log(1+x)) could potentially lead to different performance results of unsupervised analysis. Normalization is crucial to address uneven sampling depth and sparsity (high proportion of zeros) in microbiome data, like rarefying an OTU table, that is randomly sampling without replacement from each sample such that all samples have the same number of total counts (sequencing depth) ^68–71^ (http://qiime.org/scripts/single_rarefaction.html). This normalization is recommended to moderate the sensitivity of UniFrac distances to sequencing (sampling) depth ^50,72^, especially differences in the presence of rare OTUs ^48^, nonetheless it is also considered statistically improper due to the omission of data ^72^.

Another normalization was introduced in 2010 by Anders and colleagues for general sequence count data (function *varianceStabilizingTransformation* implemented in the Bioconductor DESeq2 package), that uses a Variance-Stabilization Transformation (VST) by modelling microbiome count data with Negative Binomial (NB) distribution ^69,72^.

We also provide the results with these two different normalizations, and we further confirm that the data are segregated in the HD space when pre-processed according to them, as shown in the p-value, AUC and AUPR tables in Additional file (for negative binomial, Supplementary Tables S5-6; for rarefaction, Supplementary Table S11-12). Interestingly, across all the datasets MCE decreases its performance with these pre-processing techniques, remarkably with rarefied datasets, while the other linear techniques improve in performance (Supplementary Table S6 for negative binomial; Supplementary Table S12 for rarefaction), suggesting that these adjustments linearize the datasets. Indeed, since MCE is a hierarchical technique, it needs the presence of nonlinearity to perform well. In a similar way, with these two normalizations the accuracy of MC-MCL drops down (less remarkably in the rarefaction datasets), while the performance of MCL does not increment (Supplementary Table S9 for negative binomial; Supplementary Table S14 for rarefaction). It is true that some pre-processing steps such as negative binomial tend to linearize the data but, in this manner, they can also remove important nonlinear discriminative information, as we show with the results of unsupervised analysis. Therefore, some pre-processing approaches can also cancel important nonlinear discriminant information present in the analysed data.

### Network analysis clarifies the effect of PPI-treatment on the gastric microbiota

Five major phyla have been detected in the normal gastric microbiota: *Firmicutes*, *Bacteroidetes* and *Actinobacteria* dominate the gastric fluid samples, while *Fusobacteria* and *Proteobacteria* are the most abundant phyla in gastric mucosal samples ^1^.

However, the composition and abundance of gastric microbiota may be affected by many factors, such as dietary habits, *H. pylori* infection, diseases and drugs, including PPIs ^1^.

Yet, although recent studies have highlighted the potential of these antacid drugs to affect the gastric microbiota, more knowledge needs to be gained about the association between PPI usage and the non-*H. pylori* bacteria in the stomach.

Since we wanted to investigate the effect of PPI intake on gastric microbiota in dyspepsia, we analysed: Amir4 for gastric fluid microbiota ^21^ and Paroni Sterbini et *al.* dataset ^22^ for gastric mucosal microflora, in the latter case restricting to PPI-treated *H. pylori*-negative (PPI-) and untreated *H. pylori* negative patients (HP-). In both studies, the samples from dyspeptic patients were analysed using the same next-generation sequencing technologies for direct sequencing of 16S rRNA gene amplicons, 454 Pyrosequencing.

For this purpose, we employed PC-corr algorithm, that was discussed in the Methods section named: ‘*PC-corr network*’. In brief, PC-corr discloses the discriminative network of features that are associated to a sample separation along a principal component direction. Hence, we expect that the PC-corr network of bacteria will offer a view on how the community of bacteria respond to PPI-treatment perturbation in the gastric niche (environment), in dyspeptic patients.

In Amir4 (gastric fluid), PCA revealed that gastric fluid samples were separated into two groups according to PPI treatment along PC2 and their difference is significant (p-value < 0.01) (Supplementary Figure S3). Hence, we built the PC-corr network ^62^ using the loadings of PC2 at cut-off 0.5 (Supplementary Figure S4).

Similarly for the Paroni Sterbini dataset (gastric mucosa), PCA (Supplementary Figure S5) could (significantly or close to significance) separate PPI-treated *H. pylori*-negative patients from untreated *H. pylori*-negative patients along PC2 and PC15 (p-value along PC2 = 0.014, p-value along PC15=0.054). Therefore we built the PC-corr network for both PC2 and PC15 discriminating dimension using 0.5 cut-off (Supplementary Figure S6, panel A and B).

Subsequently, to investigate how PPI is affecting the microbiota in the gastric environment, we considered the conserved network, which is obtained as the union of the two PC-corr networks (obtained for PC2 and PC15) derived from the Paroni Sterbini gastric mucosa dataset intersected with the PC-corr network derived from the Amir4 gastric fluid dataset. The resulting conserved network displays the bacteria with same trend in the two datasets, i.e. either increased or decreased with PPI-treatment, respectively in red and black colour, as emphasized by the violet circle at the centre of Figure 6. Figure 7 is the same as Figure 6 but here the nodes are coloured according to phylum-level taxonomy. The conserved network which arises at the overlap between the two PC-corr networks (union of Paroni Sterbini networks intersected with the Amir4 network) is statistically significant (p-value=1.00e-04), as a result of the statistical test based on trying to obtain the same conserved network by random resampling the bacteria in the two networks (Supplementary Figure S7), implying the difficulty of generating this intersection simply at random (since this intersection lies to the right of the critical value at the 0.05 level in the distribution of overlap). This is an important result because it confirms the robustness of the detected conserved network as a microbiota signature perturbed by PPI treatment. The top and bottom panels in Figure 6 and 7 show instead the remaining part of Amir4’s network (top panel) and of Paroni Sterbini’s network (bottom panel) that are not in the intersection, and therefore might be more specific for the gastric fluid and mucosa respectively. The PPI-perturbed conserved network is characterized by a main interconnected module with nine bacteria of four different phyla (*Bacteroidetes, Fusobacteria, Proteobacteria, Firmicutes)* that are positively associated (red edges) and by two single bacteria order without interactions (*Streptophyta, Clostridiales)*, all being increased following PPI treatment, except *Streptophyta* that is instead decreased with PPI-treatment (Fig. 6 and 7). Note that a mix between genera, phyla and order of bacteria can be found in the networks. The reason behind it is the availability of detail information regarding different bacteria. Some of the spotted bacteria (*Veillonella, Clostridiales, Campylobacter*) were already observed in previous studies. The genus *Veillonella* was found increased in relation to PPI use ^16^ in the gut microbiome and has been associated with increased susceptibility to *Clostridium difficile* infection ^73^. These Gram-negative anaerobic cocci with lactate fermenting abilities are abundant in the human microbiome and are normally found in the intestines and oral mucosa of humans ^74^. Interestingly, they favour nitrite accumulation in the stomach during nitrate reduction, promoting a carcinogenic effect ^1^. In addition, the order *Clostridiales,* that is associated to *Clostridium difficile* infection, was also seen significantly changed in the gastrointestinal tract, however Freedberg *et al.* ^4^ found it significantly decreased during PPI use, in contrast to our results. PPIs use also increases the risk of other enteric infections, apart from *C. difficile* infection, such as campylobacteriosis, as reported in ^75,76^. Moreover, half of the bacteria present in the network normally colonize the human oral cavity. Indeed, it is the main purpose of PPI treatment to increase the stomach pH, and the higher pH of treated patients is known to favour the growth of bacteria that usually reside in the mouth and esophagus and are not adapted to survive the normal gastric acidity ^6,20^. Among genera usually reported as part of the normal flora of the gastrointestinal tract, only *Veillonella* is found regularly at other sites, like the mouth ^77^. *Leptotrichia* species mostly colonize the oral cavity and they were isolated from various human infections, suggesting that they are emerging human pathogens ^78,79^. *Oribacterium* also inhabits the mouth, besides the upper respiratory tract ^80^. *Prevotella* is a genus of Gram-negative bacteria that tend to colonize the human gut, mouth and vagina, and may cause infections, mostly observed in the oral cavity (odontogenic infections) ^79^. *Porphyromonas* has been found by ^81^ as part of the salivary microbiome. Both *Prevotella* and *Porphyromonas* contribute to the formation of abscesses and soft tissue infections in various part of the body and they can cause infections, including periodontal and endodontal diseases^82^. *Capnocytophaga* are inhabitants of the oral cavity too, and these opportunistic pathogens can cause infections (both in immunocompromised and immunocompetent hosts), the severity of which depend on the immune status of the host ^83,84^. As well, *Granulicatella* are Gram-positive cocci normally found in the oral flora and are uncommon causes of infections, nevertheless they can cause infections, including bloodstream infection and infective endocarditis ^85^. Besides, the genus *Fusobacterium* inhabits the mucosal membranes of humans and all its species are parasites of humans ^86^, and some species are found in the oral cavity. The remaining bacteria (*Campylobacter, Bulleidia)* do not belong to the oral microbiota ^82^. The genus *Campylobacter* was increased in relation to PPI use and the increased abundance of these Gram-negative bacteria has the potential to cause diseases and infections in humans (most commonly diarrhoea). Due to the induced increase of pH, PPI is hypothesised to facilitate gastrointestinal infections and a study by Brophy *et al.* ^87^ reported an increased risk of *Campylobacter* infection following PPI therapy. Moreover Campylobacteriosis, mostly caused by eating undercooked foods derived from poultry or other warm-blooded animals or contact with contaminated water or ice ^88^, has been shown by the Dutch National Institute for Public Health and the Environment to noticeably increase in incidence when PPI use grows ^75^.

**Figure 6.**
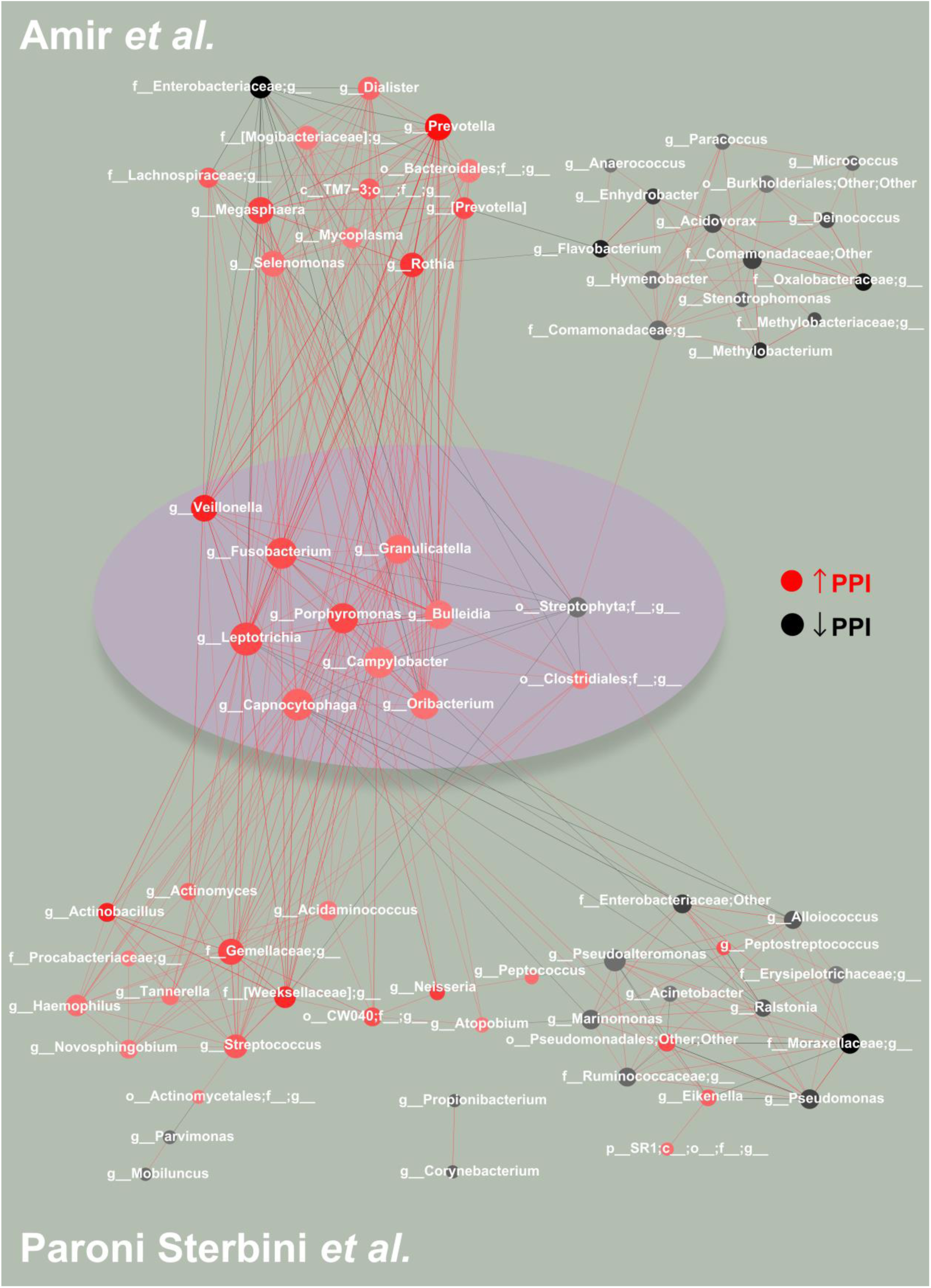
PC-corr method to unveil how PPI is affecting the microbiota in gastric environment in dyspeptic patients. **(Middle panel)** To investigate the effect of PPIs on the gastric microbiota in dyspeptic patients, we constructed the conserved PC-corr network at 0.5 cut-off, by merging the PC-corr networks obtained from the gastric mucosa (Paroni Sterbini *et al.* ^22^) and the gastric fluid (*Amir et al.* ^21^). To do so, we firstly considered the union of the two PC-corr networks obtained from the gastric tissue dataset and then we intersected it with the PC-corr network from the gastric fluid dataset. All the bacteria spotted in the conserved PC-corr network (violet circle) were found increased with PPI use. In both the two studied datasets, red nodes indicate bacteria whose abundance is increased with PPI-treatment, while black nodes indicate bacteria with lower abundance following treatment with this acid suppressing medication. The common bacteria that showed an opposite trend in the two datasets, i.e. microbial abundance increased in one dataset and decreased in the other dataset, were removed from the network. (**Top panel**) The top panel shows the obtained Amir4’s network, not in common with the Paroni Sterbini’s network. The module on the left side (except *Enterobacteriaceae)* include bacteria more abundant following PPI-treatment in Amir4’s data, while the module on the right (and *Enterobacteriacea)* is composed of decreased bacteria in abundance under PPI therapy in Amir4’s data. (**Bottom panel**) The bottom panel represents the part of Paroni Sterbini’s network (union of the two PC-corr network), that is not shared with Amir4’s one. As in the top and middle panels, the colour of the nodes represents if the bacteria display higher (red nodes) or lower abundance (black nodes) in PPI-treated samples of Paroni Sterbini’s dataset.

**Figure 7.**
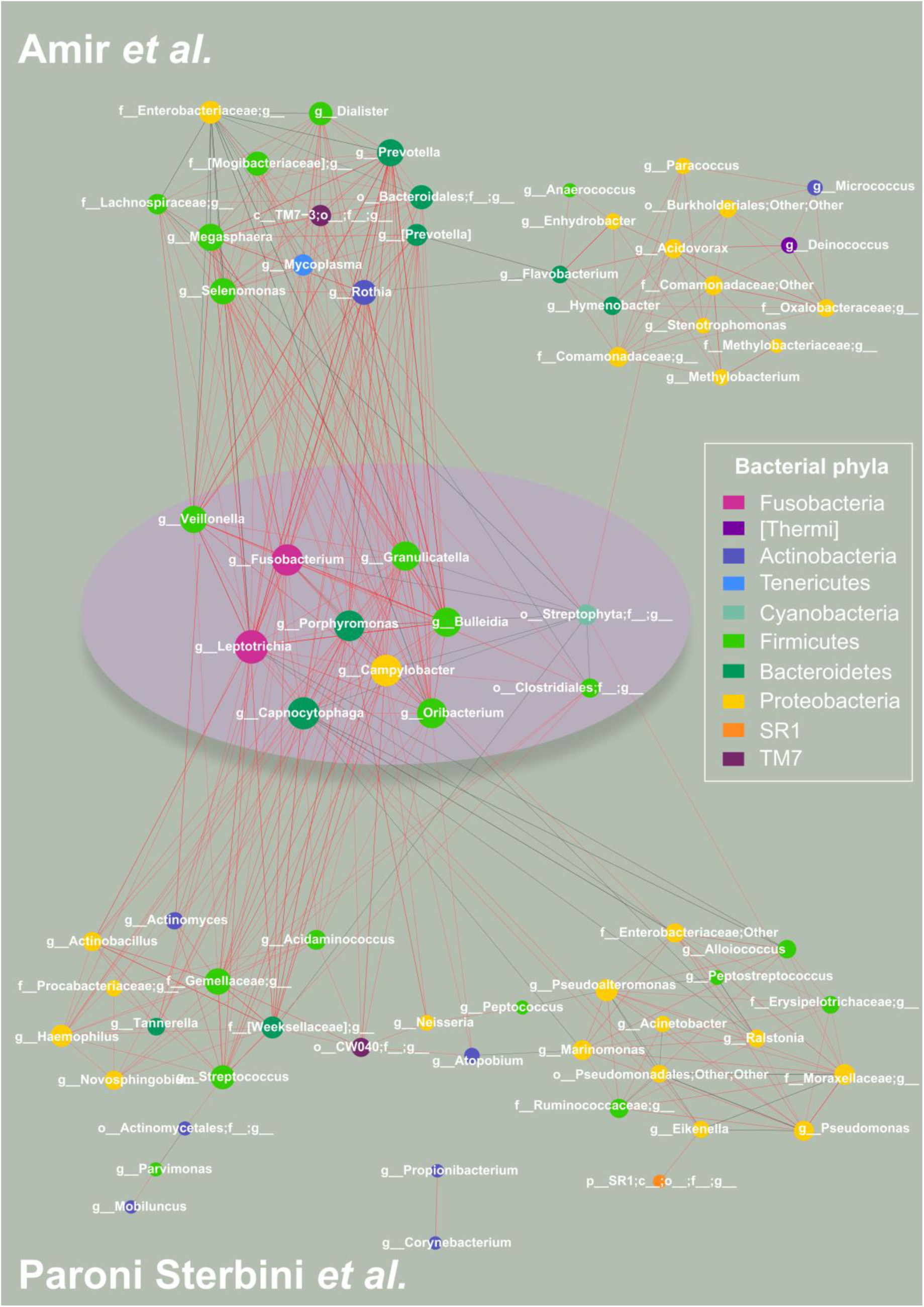
PC-corr networks to unveil how PPI is affecting the microbiota in gastric environment in dyspeptic patients, coloured according to phylum-level taxonomy. To investigate the effect of PPIs on the gastric microbiota in dyspeptic patients, we constructed the conserved PC-corr network at 0.5 cut-off, by merging the PC-corr networks obtained from the gastric mucosa (Paroni Sterbini *et al.*^22^) and the gastric fluid (*Amir et al.* ^21^). To do so, we firstly considered the union of the two PC-corr networks obtained from the gastric tissue dataset and then we intersected it with the PC-corr network from the gastric fluid dataset. All the bacteria spotted in the conserved PC-corr network (violet circle) were found increased with PPI use. (**Top panel**) The top panel shows the obtained Amir4’s network, not in common with the Paroni Sterbini’s network. The module on the left side (except *Enterobacteriaceae)* include bacteria more abundant following PPI-treatment in Amir4’s data, while the module on the right (and *Enterobacteriacea)* is composed of decreased bacteria in abundance under PPI therapy in Amir4’s data. (**Bottom panel**) The bottom panel represents the part of Paroni Sterbini’s network (union of the two PC-corr network), that is not shared with Amir4’s one. As in the top and middle panels, nodes are coloured according to bacterial phylum level.

Altogether, PC-corr approach was applied on gastric fluid and gastric mucosal datasets (in the latter case, excluding the samples positive to *H. pylori* infection) to investigate how PPI is affecting the gastric microbiota (both gastric fluid and gastric mucosal microbiota), because of PC-corr’s ability to pinpoint the combination of bacteria that play a major role in the discrimination of the samples, in this case according to PPI intake. The PC-corr conserved network identified eleven genera and order of bacteria, which belong to the phyla (*Bacteroidetes, Fusobacteria, Proteobacteria, Firmicutes)* commonly found in the stomach which, with exception of *Streptophyta*, demonstrated increased abundance following PPI treatment. Mostly all the found bacteria were not reported in previous studies, except *Veillonella*, *Clostridiales* and *Campylobacter*, but they were found as inhabitants of the oral cavity and/or possible cause of infections and diseases in humans. Hence, and in concordance to previous studies ^6,20^, these results point out that PPI treatment, by increasing the intragastric pH, favours the growth of bacteria that usually reside in the mouth and survive through the harsh acidic conditions of the stomach. Furthermore, the results suggest that PPI-associated increas of some bacterial populations may lead to infections and diseases or increase susceptibility for other bacterial infections (like *Veilonella)* or promote a carcinogenic effect (like *Veilonella).* Previous studies have highlighted that PPI intake is associated with decreased bacterial richness ^16,18,89,90^, increased risk of enteric and other infections (e.g. caused by *Salmonella, Clostridium difficile*, *Shigella, Listeria*) ^17,91^, increase in the abundance of oral and upper GI tract commensals and potential pathogenic bacteria (e.g. *Enterococcus, Streptococcus, Staphylococcus*, and *Escherichia coli)* ^16,17^ in the gut microbiota. Nevertheless, our analysis by means of PC-corr does not spot single bacteria perturbed in the gastric environment by PPI treatment, but a community of bacteria is altered in abundance by PPIs and their inter-specific bacterial interactions in the gastric niche.

Therefore our study will ground the basis for further investigations that could better clarify the effect of PPI-treatment on the human gastric microbiota and additionally verify the identified altered bacteria, as PPIs may have possible side-effects, including increased risks of different infections and diseases.

### Network analysis clarifies the effect of *H. pylori* infection on gastric mucosal microbiota

The stomach was long thought sparsely colonized by bacteria due to the gastric microbicidal acidic barrier (pH<4.0) ^92^. This view dramatically changed with the discovery of the Gram-negative bacterium *H. pylori* in the 1980’s by Warren and Marshall ^93^, that is a carcinogenic bacterial pathogen infecting the stomach of more than one*-*half of the world’s human population. This human pathogen is able to survive in the highly acidic environment within the stomach by producing cytoplasmic urease that, by catalysing the hydrolysis of urea into CO_2_ and NH_4_, produces a neutralizing ammonia cloud around it ^19,94,95^. However, most *H. pylori* avoid the acidic environment of the gastric lumen by swimming towards the mucosal cell surface (using their polar flagella and chemotaxis mechanisms) and may adhere and invade the gastric mucosal epithelial cells ^96,97^. Hence, it doesn’t represent a dominant species in gastric fluid microbiota ^98^, but was found to generally to reside in the gastric mucosae ^5,96,99^.

Persistent (chronic) infection with this Gram-negative bacterium induces changes in gastric physiology and immunology, e.g. reduced gastric acidity and parietal cell mass, perturbed nutrient availability, local innate immune responses ^100,101^, that most probably induces shift in gastric microbiota composition ^100^. Although *H. pylori* colonization usually persists in the human stomach for many decades without adverse effects, the infection of this bacteria is associated with increased risk for several diseases, including peptic ulcers, chronic gastritis, mucosa-associated lymphoid tissue lymphoma, gastric adenocarcinoma ^102,103^, and dyspepsia ^104,105^. The potential alterations induced by the *H. pylori* can in turn lead to dysbiosis and may cause aberrant proinflammatory immune responses ^106^, susceptibility to bacterial pathogens and increased risk of gastric disease, including cancer ^1,107^. However, the effect of *H. pylori* infection on overall composition of gastric microbiota at genus level and the bacterial interplay in presence of this widespread human infection remain unclear.

To investigate the influence of *H. pylori* infection on the gastric mucosal microbiota, we analysed: 1) Paroni Sterbini *et al.* ^22^ considering only PPI-untreated dyspeptic patients, either infected (HP+) or not by *H. pylori* (HP-); 2) Parsons *et al*. ^29^ restricting to PPI-untreated patients from: i) normal stomach group with no evidence of *H. pylori* infection; ii) *H. pylori* gastritis group with evidence of *H. pylori* infection. Even though the same technology is important for a comparative study, unfortunately in the literature there was no such data available like Paroni Sterbini’s one, that is 16S rRNA gene pyrosequencing data (derived from gastric mucosal microflora in dyspeptic untreated patients either positive or negative for *H. pylori*). Despite this, the two studied datasets, obtained with two different next-generation sequencing technologies for direct sequencing of 16S rRNA gene amplicons (454 Pyrosequencing for Paroni Sterbini *et al.* and Illumina MiSeq for Parsons *et al.*) ^108^, both contain community profiling of gastric mucosa-associated microbiota in PPI-untreated *H. pylori*-negative and - positive subjects. However, for the sake of clarity, we have to specify a difference: while in Paroni Sterbini’s dataset the gastric mucosal biopsy specimens were collected from patients with dyspepsia, this is not the case for Parsons’s data.

To enhance the understanding of the *H. pylori-*triggered microbial perturbation in this ecological niche, we employed again PC-corr algorithm, that is able to associate to any PCA analysis of an omic dataset, where a sample separation emerges, a network of discriminative features (for details see *‘Methods-PC-corr network’*). The analysis of the 16S rRNA sequencing data was restricted only the overlapping OTUs, excluding *Helicobacter* because our goal is to investigate its impact on the rest of the microbial network.

In Paroni Sterbini’s dataset, since PCA could significantly separate gastric mucosal biopsy samples of PPI-untreated patients according to *H. pylori*-positivity (p-value=0.01) along PC2 (Supplementary Figure S8), the PC-corr network was constructed from PC2 loadings at 0.5 cut-off (Supplementary Figure S9). Similarly, for Parsons’ dataset, since PCA (Supplementary Figure S10) could significantly separate patients from the normal stomach group with no evidence of *H. pylori* infection and PPI-untreated (Control) from *H. pylori* gastritis group positive to *H. pylori* infection and not using PPIs (HPGas) along PC1 (p-value along PC1 <0.01,), the PC-corr network was constructed from this discriminating dimension at 0.5 cut-off (Supplementary Figure 11). The obtained microbial differential networks (top panel for and bottom panel in Figure 8, coloured according to phylum level) pinpointed, from the system point of view, the bacteria affected by *H. pylori* infection in the gastric mucosa, that are precisely bacteria whose abundance is decreased in *H. pylori*-positive patients. A presumable explanation of this trend is already pointed out in literature, where the presence of *H. pylori* leads to a reduced gastric microbial diversity ^109–111^. Nevertheless, in some cases the diversity increases again, because of diverse factors that allow survival and colonization of bacteria in the stomach ^1,112^. Then, the preserved network of gastric mucosa microbiota was constructed by intersecting the two PC-corr networks obtained from Paroni Sterbini’s and Parsons’s dataset. Figure 8, middle panel, shows the conserved network (violet circle), which presents the common bacteria coloured according to phylum level and their associations. The spotted bacteria display decreased abundance with *H. pylori* infection (i.e. increased in *H. pylori-negative* subjects) in both the two 16S rRNA gene sequencing data. By performing a statistical test based on random resampling of the bacteria in the two networks, we verified that the shown bacterial conserved network is statistically significant and difficult to be generated at random (p-value=1.00e-04), because getting this intersection at random is very rare (Supplementary Figure S12). The top and bottom panels in Figure 8 show instead the remaining part of Paroni Sterbini’s network (top panel) and of Parsons’s network (bottom panel) that are not in the intersection. At the genus level, a study by Klymiuk *et al.* ^113^ identified *Actinomyces, Granulicatella, Veillonella, Fusobacterium, Neisseria, Helicobacter, Streptococcus,* and *Prevotella* as significantly different between the *H. pylori*-positive and *H. pylori*-negative gastric samples. These bacteria do not emerge in the conserved network, while they all (except *Neisseria)* appear altered (decreased) during *H. pylori* infection in the study by Parsons and colleagues (present in the bottom panel of Figure 8).

**Figure 8.**
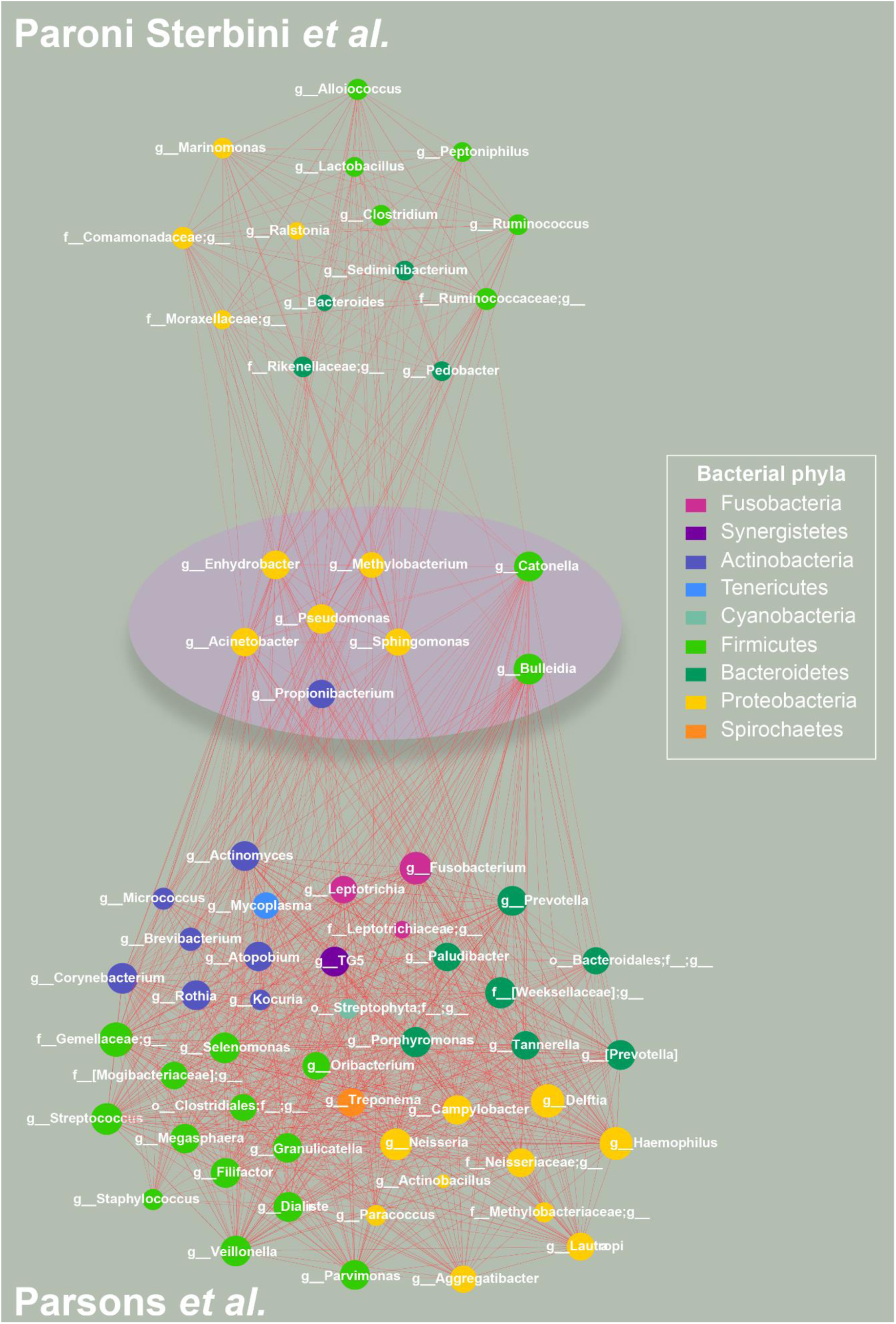
PC-corr network to investigate the effect of *H. pylori* infection on the gastric mucosal microbiota, coloured according to phylum-level taxonomy. **(Middle panel)** To investigate the effect of *H. pylori* infection on the gastric mucosal microbiota, we constructed the conserved PC-corr network at 0.5 cut-off, by intersecting the PC-corr networks obtained from Paroni Sterbini *et al.* ^22^ and *Parsons et al* ^29^ dataset. All the bacteria spotted in the conserved PC-corr network (violet circle) were found decreased in abundance with *H. pylori* infection. The common bacteria that showed an opposite trend in the two datasets, i.e. microbial abundance increased in one dataset and decreased in the other dataset, were removed from the network. (**Top panel**) The top panel show the obtained Paroni Sterbini’s network, not in common with the Parsons’s network. It contains all bacteria whose abundance is decreased in *H. pylori-*positive patients in Paroni Sterbini *et al.* dataset. (**Bottom panel**) The bottom panel represent the part of Parsons’s network that is not shared with Paroni Sterbini’s one. As in the top and middle panels, it includes bacterial communities decreased in *H. pylori-*infected patients.

Our analysis pinpoints a conserved network from two independent 16S rRNA gene sequencing data, that reveals microbial communities altered by *H. pylori* infection and their interactions in the gastric mucosa. It revealed a main core of six associated bacteria (with positive association, red edges) and two single nodes without any interaction with the main module, from three different phyla (*Proteobacteria, Firmicutes, Actinobacteria*) all resulting decreased in *H. pylori*-infected subjects (that is increased in non-infected subjects). The decreased abundance of the phyla *Firmicutes* and *Actinobacteria* in *H. pylori-*positive patients with respect to *H. pylori-*negative subjects was already shown in a previous study by Maldonado-Contreras *et al.*^114^. In addition, other studies have demonstrated an increased colonization of *Proteobacteria* in *H. pylori*-positive patients ^114,115^, while the obtained conserved PC-corr network shows that the bacteria from this phylum are instead decreased in those individuals. Among the spotted bacteria, *Methylobacterium* is a genus of facultative methylotrophic bacteria that are commonly found in diverse natural environments (such as leaf surfaces, soil, dust, and fresh water) and in hospital environment due to contaminated tap water. *Methylobacterium* species can cause health care-associated infections (mainly catheter infection), especially in immunocompromised patients ^116^. In addition, *Sphingomonas* plays a role in human health, as some of the sphingomonads (in particular *Sphingomonas paucimobilis*) are the cause of a range of mostly nosocomial, non-life-threatening infections. *Sphinhomonas* species are widely spread in nature, having been isolated from many sources, from water habitats to clinical settings ^117^, *Pseudomonas*, due to its great metabolic versatility, can also colonize different types of niches ^118^, including soil and water, in addition to plant and animal associations, and includes pathogenic species in humans ^119^. *Acinetobacter* species are instead common, free-living saprophytes found in soil, water, sewage and foods and are ubiquitous organisms in hospitals. They have been increasingly identified as a key source of infection in debilitated patients in hospitals, due to their rapid development of resistance to antimicrobials ^120^. In particular, one species, *Acinetobacter lwoffi*, can trigger gastritis, apart from *H. pylori* ^121^*. Propionibacterium,* so named for their unique ability to synthesize propionic acid by using unusual transcarboxylase enzymes ^122^, are primarily facultative pathogens and commensals of humans, living on the skin, while other members are widely employed for synthetizing vitamin B_12_, tetrapyrrole compounds, and propionic acid, as well as used as probiotics ^123^. *Catonella* is another node in the network and this bacterial genus is obligative anaerobic, non-spore-forming and non-motile, with one known species (*Catonella morbi*) from the human gingival crevice ^124,125^, that has been associated with periodontitis ^124^ and endocarditis ^126^. Besides, the bacterial genus *Enhydrobacter* so far contains a single species, *Enhydrobacter aerosaccus*, a Gram negative non-motile bacterium that is both oxidase and catalase positive and shows gas vacuoles ^127,128^. *Bulleidia*, a Gram-positive, non-spore-forming, anaerobic and non-motile genus, has one known species too (*Bulleidia extructa)*^129^.

In conclusion, by means of the PC-corr approach, we determined the combination of bacteria responsible for the difference between *H. pylori-*positive and *H. pylori*-negative gastric mucosa of untreated patients and their microbe-microbe interactions. All the bacteria, both in the conserved network and not, were decreased in *H. pylori-*infected individuals (i.e. increased in *H. pylori-*negative group). *H. pylori,* like acid suppressing medications (for the treatment of dyspepsia), alters the population structure of the gastric and intestinal microbiota ^130^ and regularly, this bacterium constitutes most of the gastric microbiota ^112^, literally depleting bacterial biodiversity. Moreover, most of the identified bacteria represent bacteria of potential health concern, as agents of diseases and infections.

## Discussion

This study indicates the necessity of including nonlinear multidimensional techniques into clinical studies based on 16S metagenomic sequencing data, since drawing a study’s conclusions by solely relying on linear techniques, such as PCA and MDS, can lead to data misinterpretation and impair the translational path from research to diagnostic. In the era of post-genomics and systems approaches, nonlinear dimension reduction and clustering by MCE and MC-MCL can offer new insights into complex clinical 16S metagenomics data, like the ones studied in this article or the presence of clinical sub-types, and serve as a valuable tool in the run towards precision medicine. Moreover, this study shows how it is possible to complement multivariate analysis by means of network analysis employing PC-corr algorithm, that accounts for the bacteria responsible for the sample discrimination and their co-occurrence relationships. Precisely, from the system point of view the obtained microbial differential networks pinpointed marked bacteria-bacteria interactions and modules affected by PPI treatment in the gastric environment in dyspepsia and by *H. pylori* infection in the gastric mucosa. We suggest that our findings can be an important starting point to design new therapies that consider not only *H. pylori* infection but also the directly associated microbial alterations as well as the indirect alterations due to the drugs used for *H. pylori* eradication such as PPI.

## List of abbreviations

LDA: Linear Discriminant Analysis
MC: Minimum Curvilinearity
MCE: Minimum Curvilinear Embedding
MCL: Markov Clustering
MC-MCL: Minimum Curvilinear Markov Clustering
MDS: Multidimensional Scaling
MDSbc: Multidimensional Scaling with Bray-Curtis dissimilarity
MDSwUF: Multidimensional Scaling with weighted UniFrac distance
MST: minimum spanning tree
ncMCE: non-centred Minimum Curvilinear Embedding
NMDS: non-metric (Sammon criterion) Multidimensional Scaling
PC: Principal Component
PCA: Principal Component Analysis
PCoA: Principal Coordinate Analysis
PPI: Proton Pump Inhibitor
PSI: Projection-based separability index
SVD: Singular Value Decomposition

## Declarations

### Ethics approval and consent to participate

Not applicable, because the used datasets have been generated by previous biomedical studies, for which ethics approvals and consents were formerly collected.

### Consent for publication

Not applicable

### Availability of data and materials

Not applicable.

### Authors’ contributions

CVC developed Minimum Curvilinearity (MCE), Minimum Curvilinear Markov Clustering (MC-MCL) and the Projection-based Separability Index (PSI). CVC conceived all the study and the data analysis workflow with feedbacks from MiSc and SWG. SC, CD and AP performed the computational analysis of the data and realized the figures under the CVC guidance. SC, CD, AP together with CVC wrote the manuscript with valuable suggestions of PS. FPS, LM, GC, GI, BP, MaSa, GG and AG provided data and knowledge about the Paroni Sterbini *et al.* data cohort. BNP, UZI and MP provided data and knowledge about the Parsons *et al.* data cohort. All authors discussed the results and revised the manuscript.

## Acknowledgements

Not applicable

