## Supplementary material for "Machine learning pattern recognition and differential network analysis of gastric microbiome in the presence of proton pump inhibitor treatment or *Helicobacter pylori* infection": suppl info

### Additional file 1: Supplementary figures and tables

#### Table of contents

**Supplementary Figure S1:** LDA analysis of gastric biopsies dataset (Paroni Sterbini *et al.*).

**Supplementary Figure S2:** MCE on gastric biopsies dataset (Paroni Sterbini *et al.*), restricted to PPI-treated patients.

**Supplementary Table S5:** Average p-value, AUC and AUPR best results with standard error on the original datasets, when applying Leave-one-out-cross-validation (LOOCV).

**Supplementary Table S6:** p-value, AUC and AUPR results on the datasets after approximation to the negative binomial distribution.

**Supplementary Table S8:** Rank performance on the datasets after approximation to the negative binomial distribution.

**Supplementary Table S9:** Clustering results on the datasets after approximation to the negative binomial distribution.

**Supplementary Table S11:** p-value, AUC and AUPR results on the rarefied datasets.

**Supplementary Table S13:** Rank performance on the rarefied datasets.

**Supplementary Table S14:** Clustering results on the rarefied datasets.

**Supplementary Figure S3:** PCA analysis reveals separation related to PPI-treatment in gastric fluid.

**Supplementary Figure S4:** PC-corr network to investigate the effect of PPI treatment on gastric fluid.

**Supplementary Figure S5:** PCA analysis reveals separation related to PPI-treatment in gastric mucosa, in the patients negative to *H. pylori* test.

**Supplementary Figure S6:** PC-corr network to investigate the effect of PPI treatment on gastric mucosa.

**Supplementary Figure S7**: The overlap between Amir *et al.* and Paroni Sterbini *et al.* networks, related to PPI treatment in dyspepsia, is statistically significant and hence it cannot be generated by a random process.

**Supplementary Figure S8:** In Paroni Sterbini *et al.* dataset, PCA analysis reveals separation related to *H. pylori* infection in gastric tissue, in the PPI-untreated patients.

**Supplementary Figure S9:** PC-corr network to investigate the effect of *H. pylori* infection on gastric mucosal microbiota in Paroni Sterbini *et al.* data.

**Supplementary Figure S10:** In Parsons *et al. dataset,* PCA analysis can significantly discriminate gastric mucosal biopsy specimens according to *H. pylori*-positivity.

**Supplementary Figure S11:** PC-corr network to investigate the effect of *H. pylori* infection on gastric mucosal microbiota in Parsons *et al.* data.

**Supplementary Figure S12:** The overlap between Paroni Sterbini *et al.* and Parsons *et al.* networks, exemplifying the effect of *H. pylori* infection on gastric mucosal microbiota, is statistically different from a random overlap.

**Supplementary Tables S1, S2, S3, S4, S7, S10, S12, S15** are Excel spreadsheets and uploaded as separate files.

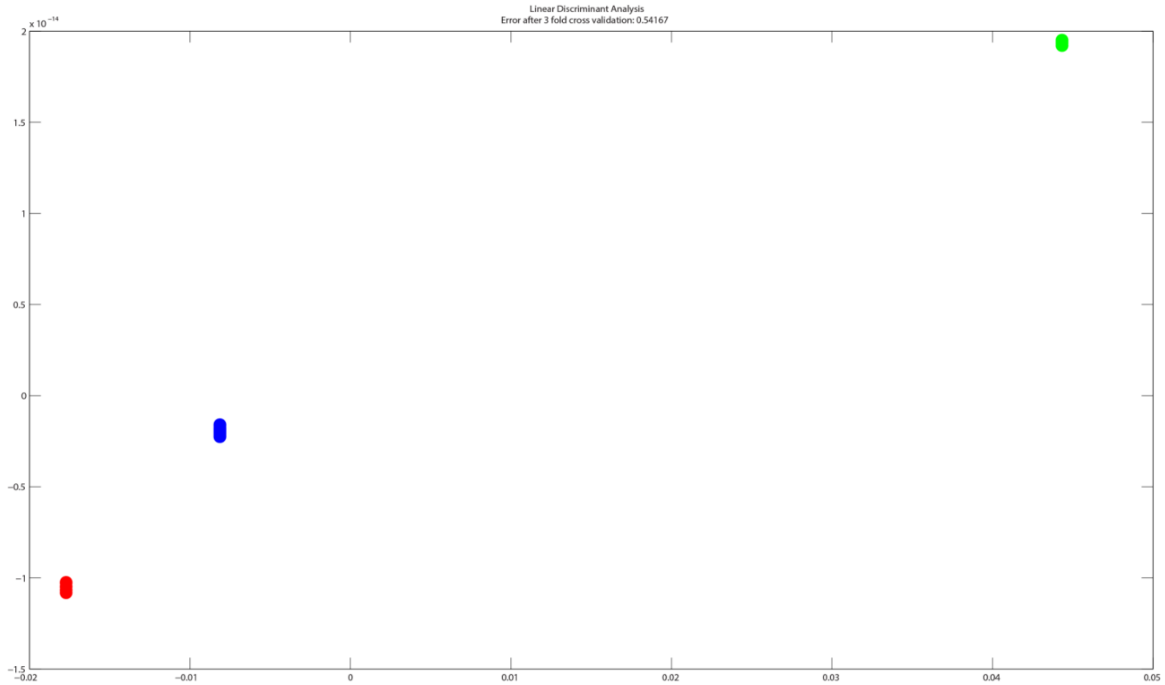


**Figure S1. LDA analysis of gastric biopsies dataset (Paroni Sterbini *et al.*).** The cross-validation test showed that this constrained technique could re-assign samples to their three groups with 54% of error, confirming its statistical invalidity for the small size dataset problem.

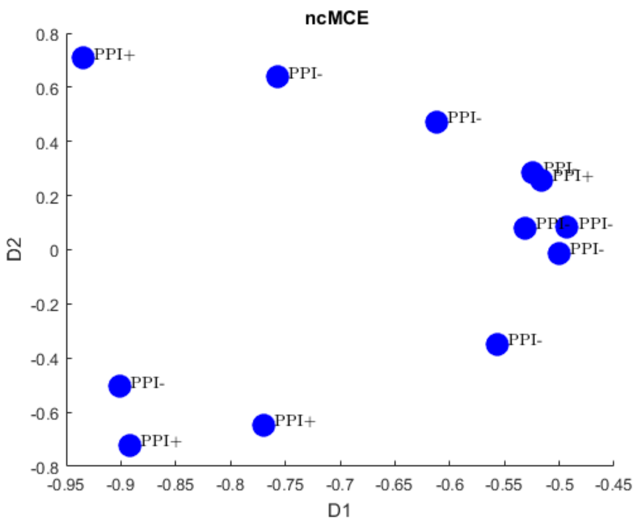


**Figure S2. MCE on gastric biopsies dataset (Paroni Sterbini *et al.*), restricted to PPI-treated patients.**

The plot shows the MCE result (ncMCE) on the gastric biopsies dataset, restricted to PPI-treated patients.

There is no internal separation of the samples related to *H. pylori* infection.

**p-value**

| Method | Paroni Sterbini | Amir3 | Amir4 |
|---|---|---|---|
| **HD** | 0.0057±0.0003 | 0.0019±0.0003 | 0.0007±0.0001 |
| **MCE** | 0.0086±0.0011 | 0.0142±0.0032 | 0.0090±0.0017 |
| **MDSwUF** | 0.0168±0.0018 | 0.0003±0 | 0.0123±0.0019 |
| **PCA** | 0.0107±0.0010 | 0.0131±0.0018 | 0.0185±0.0027 |
| **NMDS** | 0.0109±0.0010 | 0.0206±0.0028 | 0.0262±0.0038 |
| **MDSbc** | 0.0266±0.0019 | 0.0192±0.0022 | 0.0238±0.0042 |

**AUC**

| Method | Paroni Sterbini | Amir3 | Amir4 |
|---|---|---|---|
| **HD** | 0.9446±0.0018 | 0.9554±0.0049 | 0.9844±0.0036 |
| **MCE** | 0.9344±0.0036 | 0.8873±0.0102 | 0.9040±0.0085 |
| **MDSwUF** | 0.9171±0.0019 | 1±0 | 0.8884±0.0077 |
| **PCA** | 0.9267±0.0022 | 0.8850±0.0078 | 0.8683±0.0073 |
| **NMDS** | 0.9262±0.0021 | 0.8627±0.0071 | 0.8504±0.0073 |
| **MDSbc** | 0.8994±0.0026 | 0.8661±0.0075 | 0.8594±0.0093 |

| AUPR | | | |
|---|---|---|---|
| **Method** | **Paroni Sterbini** | **Amir3** | **Amir4** |
| **HD** | 0.9710±0.0011 | 0.9581±0.0054 | 0.9853±0.0037 |
| **MCE** | 0.9699±0.0017 | 0.9107±0.0093 | 0.9176±0.0073 |
| **MDSwUF** | 0.9440±0.0019 | 1±0 | 0.9071±0.0065 |
| **PCA** | 0.9565±0.0016 | 0.8953±0.0076 | 0.8894±0.0071 |
| **NMDS** | 0.9549±0.0015 | 0.8818±0.0080 | 0.8800±0.0071 |
| **MDSbc** | 0.9247±0.0026 | 0.8898±0.0066 | 0.8962±0.0073 |

**Table S5. Average p-value, AUC and AUPR best results with standard error on the original datasets, when applying Leave-one-out-cross-validation (LOOCV).** The table shows the average best results of index for sample separation in the space of the first two dimensions of embedding based on the well-known metrics Mann-Whitney (MW) p-value, Area Under the ROC-Curve (AUC) and Area Under the Precision-Recall curve (AUPR) (regardless of the normalization and type of correlation, and the type of MCE) performed in each of the three different datasets presented in the article (Paroni Sterbini, Amir3 and Amir4), with the respective standard error. This was done by applying Leave-one-out-cross-validation (LOOCV) where for each dataset the average results and the respective standard error are obtained from the best results of each leave-one-out cross-validation of the dataset (where one sample per time was removed from the dataset and then put back after the analysis was done).

| p-value | | | | |
|---|---|---|---|---|
| **Method** | **Paroni Sterbini** | **Amir3** | **Amir4** | **mean** |
| **HD** | **0.002293** | **0.000155** | **0.001865** | **0.001438** |
| **MDSwUF** | **0.011043** | **0.000311** | **0.010412** | **0.007255** |
| **MCE** | **0.010561** | **0.001088** | **0.014763** | **0.008804** |
| **PCA** | **0.001823** | **0.004662** | **0.028127** | **0.011537** |
| **MDSbc** | **0.017314** | **0.006993** | **0.020668** | **0.014992** |
| **NMDS** | **0.007221** | **0.006993** | **0.037918** | **0.017377** |

**AUC**

| Method | Paroni Sterbini | Amir3 | Amir4 | mean |
|---|---|---|---|---|
| **HD** | 0.961905 | 1 | 0.9375 | 0.966468 |
| **MDSwUF** | 0.919841 | 0.984375 | 0.875 | 0.926405 |
| **MCE** | 0.907937 | 0.953125 | 0.859375 | 0.906812 |
| **PCA** | 0.968254 | 0.90625 | 0.828125 | 0.900876 |
| **MDSbc** | 0.894444 | 0.890625 | 0.84375 | 0.876273 |
| **NMDS** | 0.92381 | 0.890625 | 0.8125 | 0.875645 |

**AUPR**

| Method | Sterbini | Amir3 | Amir4 | mean |
|---|---|---|---|---|
| **HD** | 0.981597 | 1 | 0.947463 | 0.976353 |
| **MDSwUF** | 0.946463 | 0.985243 | 0.901061 | 0.944255 |
| **MCE** | 0.960936 | 0.947917 | 0.857434 | 0.922095 |
| **PCA** | 0.985152 | 0.890625 | 0.864441 | 0.913406 |
| **MDSbc** | 0.948939 | 0.890625 | 0.876854 | 0.905473 |
| **NMDS** | 0.963295 | 0.890625 | 0.855934 | 0.903285 |

*Note: all the p-values, AUC and AUPR can be found in Supplementary Table S7*

**Table S6. p-value, AUC and AUPR results on the datasets after approximation to the negative binomial distribution.** The table shows the best results of index for sample separation in the space of the first two dimensions of embedding, based on the well-known metrics Mann-Whitney (MW) p-value, Area Under the ROC-Curve (AUC) and Area Under the Precision-Recall curve (AUPR) (regardless of the normalization and type of correlation, and the type of MCE) performed in each of the three different datasets presented in the article (Paroni Sterbini, Amir3 and Amir4), after approximation to the negative binomial distribution, and the mean performance (mean of the lowest p-values, AUC and AUPR per method) across all the datasets. Bold values indicate a significant (<0.05) p-value.

Results are ordered from the best (top) to the worst (bottom) method. For Paroni Sterbini dataset, we show the results for three different labels (PPI-treated, untreated HP+ and untreated HP-). Instead, for Amir datasets, the p-values were computed for two groups, i.e. presence or absence of PPI treatment.

**p-value**

| Method | Paroni Sterbini | Amir3 | Amir4 | mean |
|---|---|---|---|---|
| **HD** | 2 | 1 | 1 | 1.333333 |
| **MDSwUF** | 5 | 2 | 2 | 3 |
| **MCE** | 4 | 3 | 3 | 3.333333 |
| **PCA** | 1 | 4 | 5 | 3.333333 |
| **MDSbc** | 6 | 5 | 4 | 5 |
| **NMDS** | 3 | 5 | 6 | 4.666667 |

**AUC**

| Method | Paroni Sterbini | Amir3 | Amir4 | mean |
|---|---|---|---|---|
| **HD** | 2 | 1 | 1 | 1.333333 |
| **MDSwUF** | 4 | 2 | 2 | 2.666667 |
| **MCE** | 5 | 3 | 3 | 3.666667 |
| **PCA** | 1 | 4 | 5 | 3.333333 |
| **MDSbc** | 6 | 5 | 4 | 5 |
| **NMDS** | 3 | 5 | 6 | 4.666667 |

**AUPR**

| Method | Paroni Sterbini | Amir3 | Amir4 | mean |
|---|---|---|---|---|
| **HD** | 2 | 1 | 1 | 1.333333 |
| **PCA** | 1 | 4 | 4 | 3 |
| **MDSwUF** | 6 | 2 | 2 | 3.333333 |
| **MCE** | 4 | 3 | 5 | 4 |
| **MDSbc** | 5 | 4 | 3 | 4 |
| **NMDS** | 3 | 4 | 6 | 4.333333 |

**Table S8. Rank performance on the datasets after approximation to the negative binomial distribution.** The table shows the rank performance of each method for each in index for sample separation in the space of the first two dimensions of embedding, based on p-value, AUC or AUPR, for the three datasets presented in the article (Paroni Sterbini, Amir3 and Amir4), after approximation to the negative binomial distribution. Each rank is related with the results obtained in Table S6. The results are ordered by the mean performance (fourth column) from the best (top) to the worst (bottom) method.

| Accuracy | Paroni Sterbini et al. (gastric biopsies) | Amir3 et al. (esophageal biopsies) | Amir4 et al. (gastric fluid) | Mean performance |
|---|---|---|---|---|
| **MC-MCL** | 0.54 (0.58) | 0.75 | 0.63 | 0.64 |
| **MCL** | 0.58 (0) | 0.69 | 0.75 | 0.67 |

*Note: all the p-values, AUC and AUPR can be found in Supplementary Table S10*

**Table S9. Clustering results on the datasets after approximation to the negative binomial distribution.** The table shows the best results of clustering (highest accuracies, regardless of the normalization and type of correlation) by Markov Clustering (MCL) and Minimum Curvilinear Markov Clustering, with square rooting the distances, in each of the three different datasets presented in the article (Paroni Sterbini, Amir3 and Amir4 datasets), after approximation to the negative binomial distribution, and the mean performance (mean of the highest accuracies) across all the datasets.

For Paroni Sterbini dataset, we show the results for three clusters (PPI-treated, untreated HP+ and untreated HP-) and in brackets the results for four clusters (PPI-treated HP+, PPI-treated HP-, untreated HP+ and untreated HP-). Instead for Amir datasets, the accuracies were computed for two groups, related to presence or absence of PPI treatment.

**p-value**

| Method | Paroni Sterbini | Amir3 | Amir4 | mean |
|---|---|---|---|---|
| **HD** | **0.002293** | **0.000155** | **0.004662** | **0.00237** |
| **MDSwUF** | **0.008965** | **0.000155** | **0.006993** | **0.005371** |
| **PCA** | **0.003991** | **0.014763** | **0.020668** | **0.013141** |
| **NMDS** | **0.005922** | **0.014763** | **0.020668** | **0.013785** |
| **MCE** | **0.025603** | **0.020668** | **0.002953** | **0.016408** |
| **MDSbc** | **0.011336** | **0.020668** | **0.020668** | **0.017557** |

**AUC**

| Method | Paroni Sterbini | Amir3 | Amir4 | mean |
|---|---|---|---|---|
| **HD** | 0.961905 | 1 | 0.90625 | 0.956052 |

| | | | | |
|---|---|---|---|---|
| **MDSwUF** | 0.92381 | 1 | 0.890625 | 0.938145 |
| **PCA** | 0.939683 | 0.859375 | 0.84375 | 0.880936 |
| **MCE** | 0.871429 | 0.84375 | 0.921875 | 0.879018 |
| **NMDS** | 0.931746 | 0.859375 | 0.84375 | 0.87829 |
| **MDSbc** | 0.910317 | 0.84375 | 0.84375 | 0.865939 |

**AUPR**

| Method | Paroni Sterbini | Amir3 | Amir4 | mean |
|---|---|---|---|---|
| **HD** | 0.983254 | 1 | 0.924621 | 0.969292 |
| **MDSwUF** | 0.947884 | 1 | 0.90758 | 0.951822 |
| **MDSbc** | 0.957403 | 0.876854 | 0.872585 | 0.902281 |
| **MCE** | 0.909312 | 0.864007 | 0.914608 | 0.895976 |
| **PCA** | 0.972905 | 0.850167 | 0.860457 | 0.89451 |
| **NMDS** | 0.961731 | 0.859375 | 0.860457 | 0.893854 |

*Note: all the P-values, AUC and AUPR can be found in Supplementary Table 12*

**Table S11. p-value, AUC and AUPR results on the rarefied datasets.** The table shows the best results of index for sample separation in the space of the first two dimensions of embedding, based on the well-known metrics Mann-Whitney (MW) p-value, Area Under the ROC-Curve (AUC) and Area Under the Precision-Recall curve (AUPR) (regardless of the normalization and type of correlation, and the type of MCE) performed in each of the three different datasets presented in the article (Paroni Sterbini, Amir3 and Amir4), after being rarefied, and the mean performance (mean of the lowest p-values, AUC and AUPR per method) across all the datasets. Bold values represent a significant (<0.05) p-value.

Results are ordered from the best (top) to the worst (bottom) method. For Paroni Sterbini dataset, we show the results for three different labels (PPI-treated, untreated HP+ and untreated HP-). Instead, for Amir datasets, the p-values were computed for two groups, i.e. presence or absence of PPI treatment.

**p-value**

| Method | Paroni Sterbini | Amir3 | Amir4 | mean |
|---|---|---|---|---|
| **HD** | 1 | 1 | 2 | 1.333333 |
| **MDSwUF** | 4 | 1 | 3 | 2.666667 |

| | | | | |
|---|---|---|---|---|
| **PCA** | 2 | 2 | 4 | 2.666667 |
| **NMDS** | 3 | 2 | 4 | 3 |
| **MCE** | 6 | 3 | 1 | 3.333333 |
| **MDSbc** | 5 | 3 | 4 | 4 |

**AUC**

| Method | Paroni Sterbini | Amir3 | Amir4 | mean |
|---|---|---|---|---|
| **HD** | 1 | 1 | 2 | 1.333333 |
| **NMDS** | 3 | 3 | 1 | 2.333333 |
| **MDSwUF** | 4 | 1 | 3 | 2.666667 |
| **PCA** | 2 | 2 | 4 | 2.666667 |
| **MCE** | 6 | 2 | 4 | 4 |
| **MDSbc** | 5 | 3 | 4 | 4 |

**AUPR**

| Method | Paroni Sterbini | Amir3 | Amir4 | mean |
|---|---|---|---|---|
| **HD** | 1 | 1 | 1 | 1 |
| **MDSwUF** | 5 | 1 | 3 | 3 |
| **MDSbc** | 4 | 2 | 4 | 3.333333 |
| **MCE** | 6 | 3 | 2 | 3.666667 |
| **PCA** | 2 | 5 | 5 | 4 |
| **NMDS** | 3 | 4 | 5 | 4 |

**Table S13. Rank performance on the rarefied datasets.** The table shows the rank performance of each method for each in index for sample separation in the space of the first two dimensions of embedding, based on p-value, AUC or AUPR, for the three datasets presented in the article (Paroni Sterbini, Amir3 and Amir4), after being rarefied. Each rank is related with the results obtained in Table S11. The results are ordered by the mean performance (fourth column) from the best (top) to the worst (bottom) method.

| Accuracy | Paroni Sterbini et al. (gastric biopsies) | Amir3 et al. (esophageal biopsies) | Amir4 et al. (gastric fluid) | Mean performance |
|---|---|---|---|---|
| **MC-MCL** | 0.58 (0.54) | 0.81 | 0.75 | 0.71 |
| **MCL** | 0.58 (0.42) | 0.69 | 0.75 | 0.67 |

*Note: all the p-values, AUC and AUPR can be found in Supplementary Table S15*

**Table S14. Clustering results on the rarefied datasets.** The table shows the best results of clustering (highest accuracies, regardless of the normalization and type of correlation) by Markov Clustering (MCL) and Minimum Curvilinear Markov Clustering, with square rooting the distances, in each of the three different datasets presented in the article (Paroni Sterbini, Amir3 and Amir4 datasets), after being rarefied, and the mean performance (mean of the highest accuracies) across all the datasets.

For Paroni Sterbini dataset, we show the results for three clusters (PPI-treated, untreated HP+ and untreated HP-) and in brackets the results for four clusters (PPI-treated HP+, PPI-treated HP-, untreated HP+ and untreated HP-). Instead for Amir datasets, the accuracies were computed for two groups, related to presence or absence of PPI treatment.

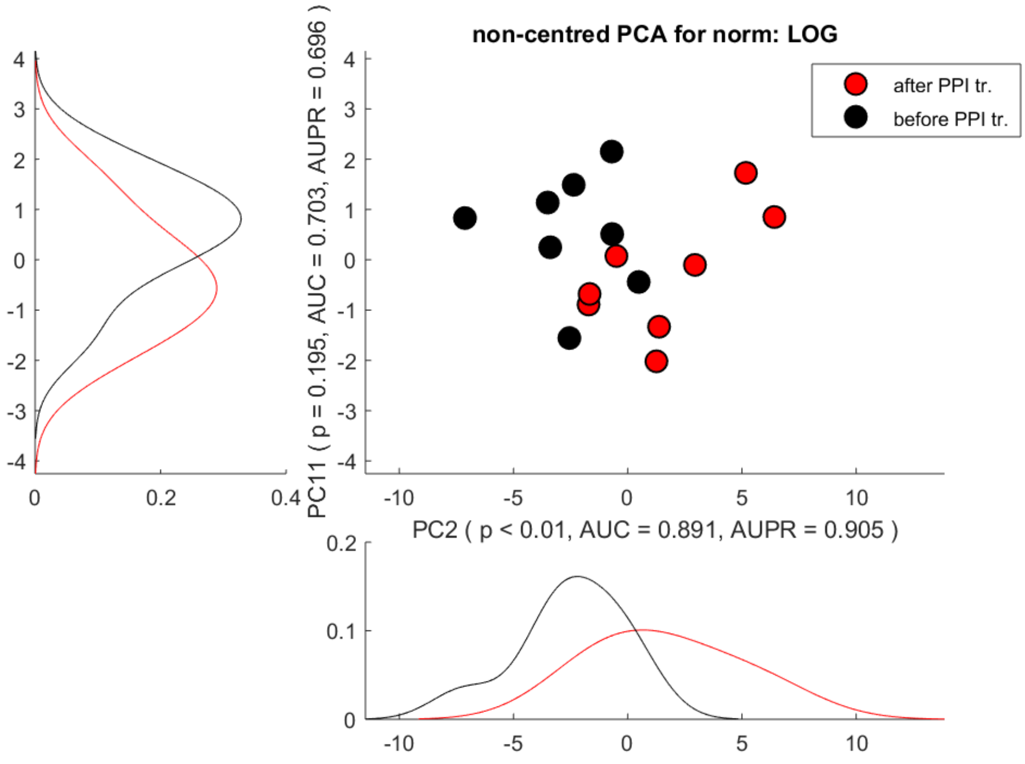


**Figure S3. PCA analysis reveals separation related to PPI-treatment in gastric fluid.** Gastric fluid samples before (black dots) and after PPI treatment (red dots) are significantly separated along PC2 (p-value <0.01).

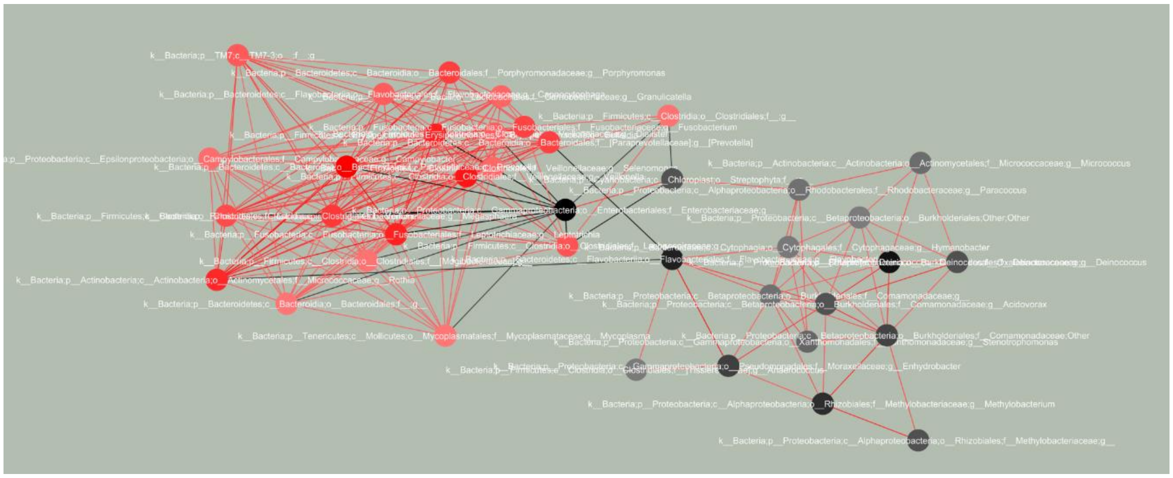


**Figure S4. PC-corr network to investigate the effect of PPI treatment on gastric fluid.** The PC-corr network was constructed at cut-off 0.5 according to the loadings of PC2, since PCA could significantly (p < 0.01) separate gastric fluid samples in individuals before and after PPI treatment (Fig. S3), therefore reflects discriminative network modules related to PPI treatment. Red nodes indicate higher bacterial abundance following PPI treatment (↑after PPI tr.), while black nodes indicate higher bacterial abundance before PPI treatment (↑before PPI tr.).

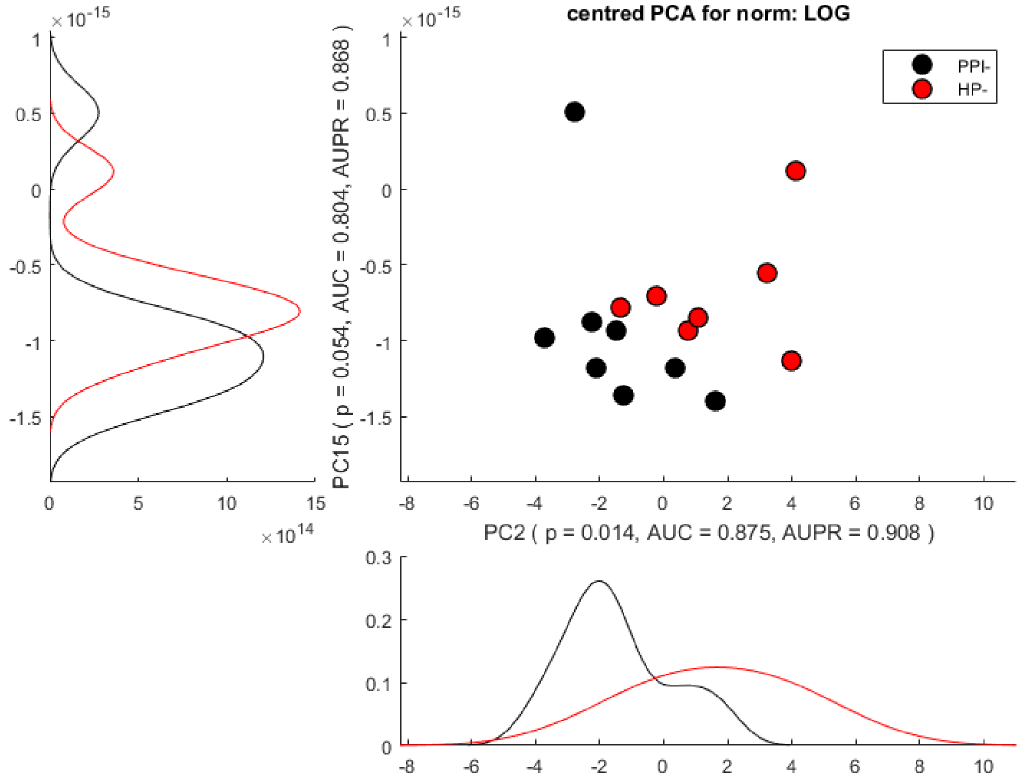


**Figure S5. PCA analysis reveals separation related to PPI-treatment in gastric mucosa, in the patients negative to *H. pylori* test.** Linear dimensionality reduction by PCA separates the gastric biopsy samples of PPI-treated *H. pylori*-negative patients (PPI -) (black dots) from the ones of untreated *H. pylori*-negative patients (HP-) (red dots) along PC2 and PC15 (significant p-value along PC2 = 0.014, p-value close to significance along PC15=0.054).

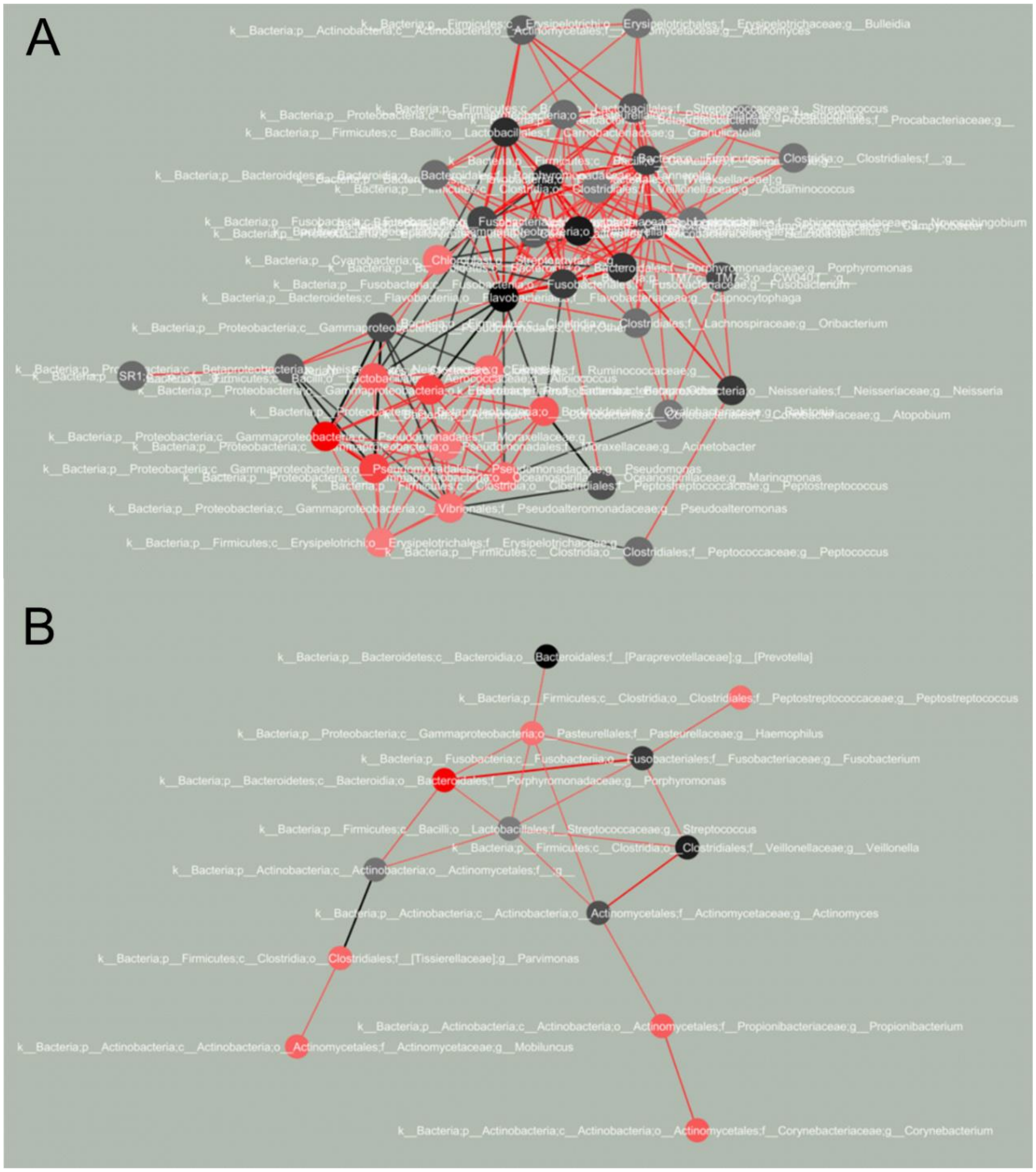


**Figure S6. PC-corr network to investigate the effect of PPI treatment on gastric mucosa.** The PC-corr network was constructed at cut-off 0.5 according to the loadings of PC2 (panel A) and PC15 (panel B), since PCA could significantly/closely to significance (p-value along PC2=0.014, p-value along PC15=0.054) separate PPI-treated *H. pylori*-negative patients from *untreated H. pylori*-negative patients (Fig. S5). Therefore, the discriminative network modules are related to PPI treatment (without *H. pylori* infection). Red nodes indicate higher bacterial abundance in untreated *H. pylori* negative patients (↑HP-), while black nodes indicate higher bacterial abundance in treated *H. pylori* negative patients (↑PPI-).

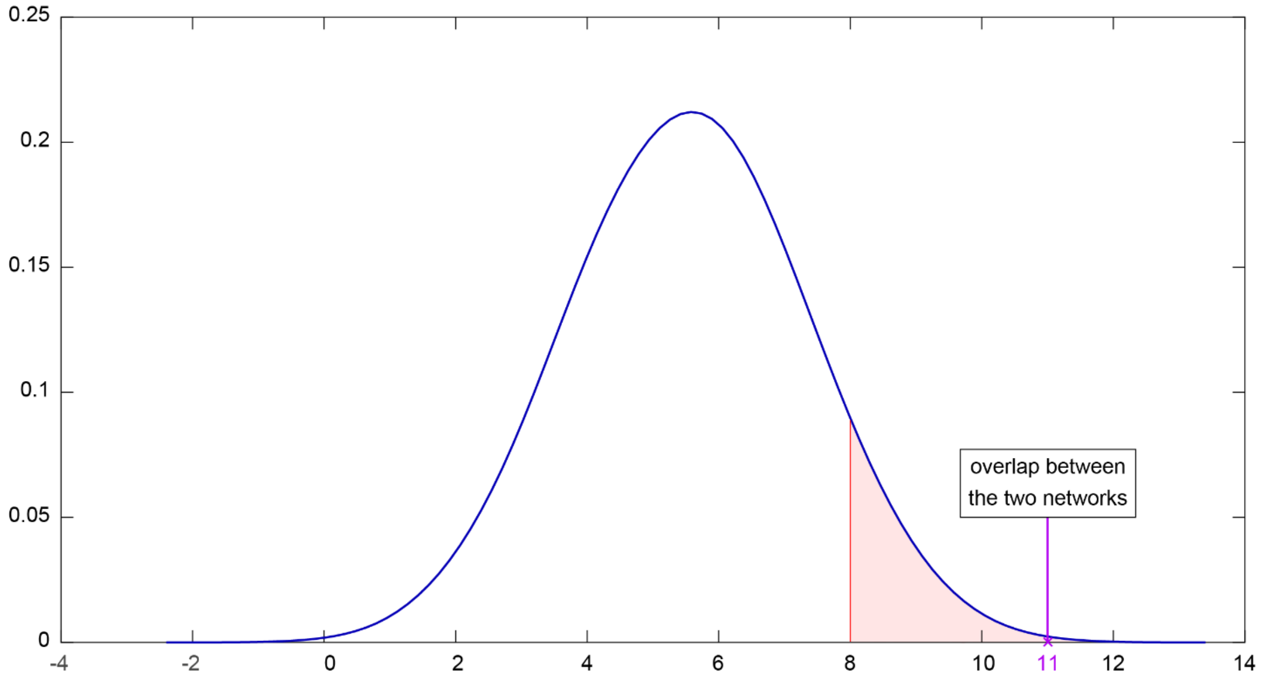


**Figure S7. The overlap between Amir *et al.* and Paroni Sterbini *et al.* networks, related to PPI treatment in dyspepsia, is statistically significant and hence it cannot be generated by a random process.** To verify that the overlap between the two networks (violet circle in Figure 6 and 7) is statistically different from a random overlap, we performed a statistical test based on random resampling of the bacteria in the two networks (repeated 10,000 times). The distribution of the resulting overlap is shown in the above figure, where the red line denotes the 95% percentile hence the tail (red area) on its right sides includes all the overlaps that are significantly different (higher) from random-overlap (p-value<0.05). The overlap between Amir *et al.* and Paroni Sterbini *et al.* networks related to PPI treatment discrimination in dyspepsia (11 bacteria, violet line) is statistically significant (p-value=1.00e-04), because getting at random this intersection is rare.

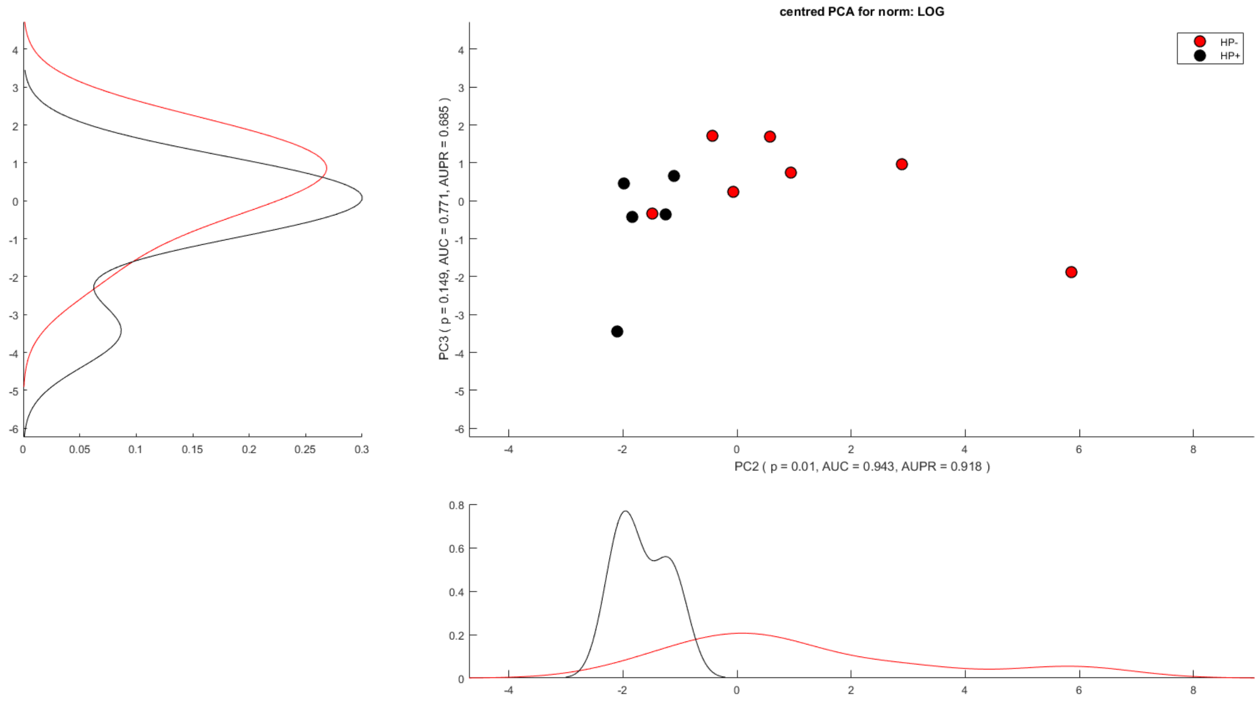


**Figure S8. In Paroni Sterbini *et al.* dataset, PCA analysis reveals separation related to *H. pylori* infection in gastric tissue, in the PPI-untreated patients.** Gastric mucosal biopsy sample from *H. pylori*-positive (HP+) (black dots) and *H. pylori*-negative (HP-) (red dots) PPI-untreated patients were seen to separate along the second principal component (PC2) (p-value=0.01).

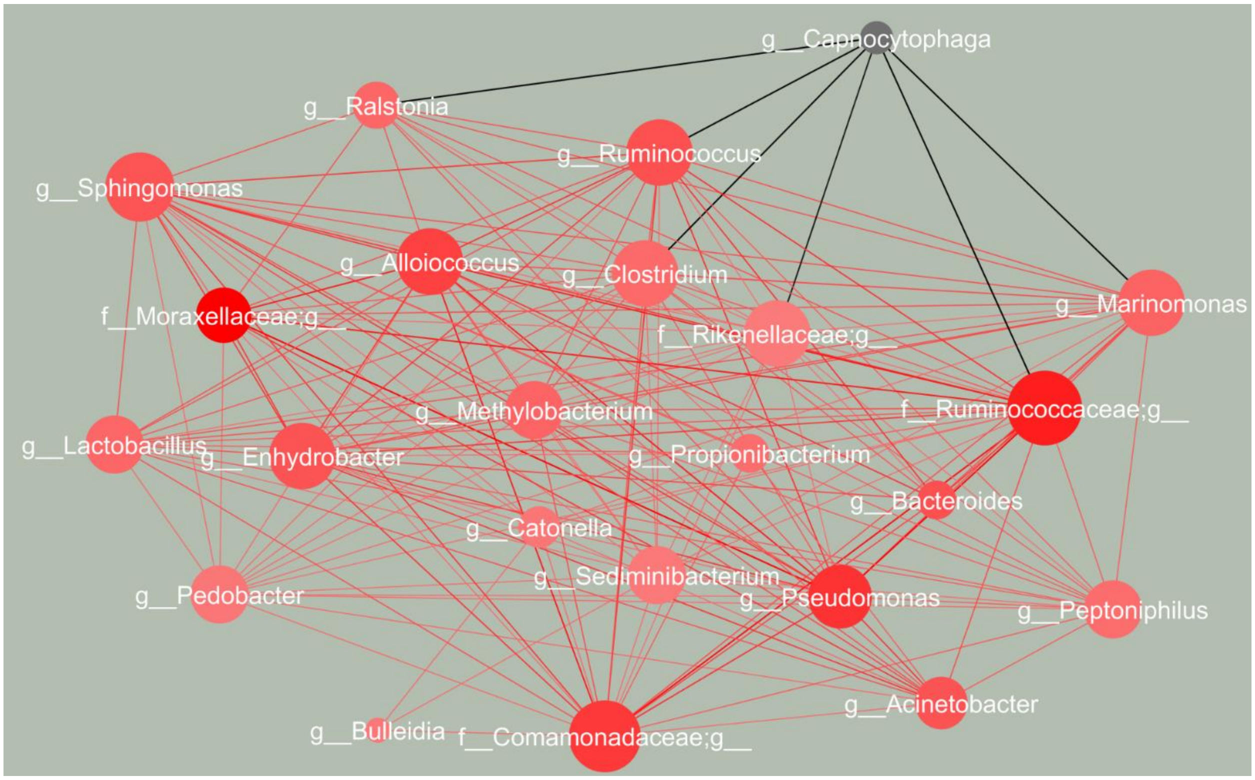


**Figure S9. PC-corr network to investigate the effect of *H. pylori* infection on gastric mucosa in Paroni Sterbini *et al.* data.** The PC-corr network was constructed at cut-off 0.5 according to the loadings of PC2, since PCA could significantly (p-value=0.01) separate PPI-untreated *H. pylori*-negative patients from PPI-untreated *H. pylori*-positive patients (Fig. S7), therefore reflects discriminative network modules related to *H. pylori* infection in gastric mucosa. Red nodes indicate higher bacterial abundance in untreated *H. pylori*-negative patients (↑HP-), while black nodes indicate bacterial abundance in untreated *H. pylori*-positive patients (↑HP+).

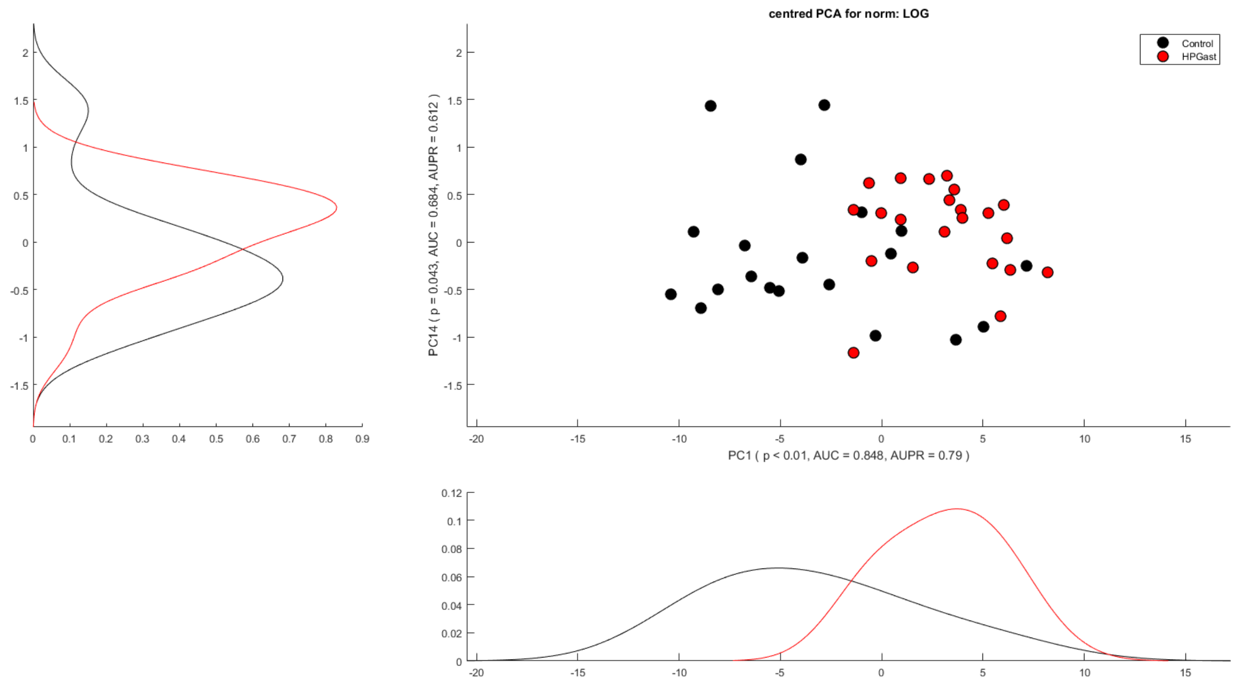


**Figure S10. In Parsons *et al. dataset,* PCA analysis can significantly discriminate gastric mucosal biopsy specimens according to *H. pylori*-positivity.** Gastric mucosal biopsy sample from normal stomach group with no evidence of *H. pylori* infection and PPI-untreated (Control, black dots) and *H. pylori* gastritis group positive to *H. pylori* infection and not using PPIs (HPGas, red dots) were seen to separate along the first principal component (PC1) (p-value<0.01).

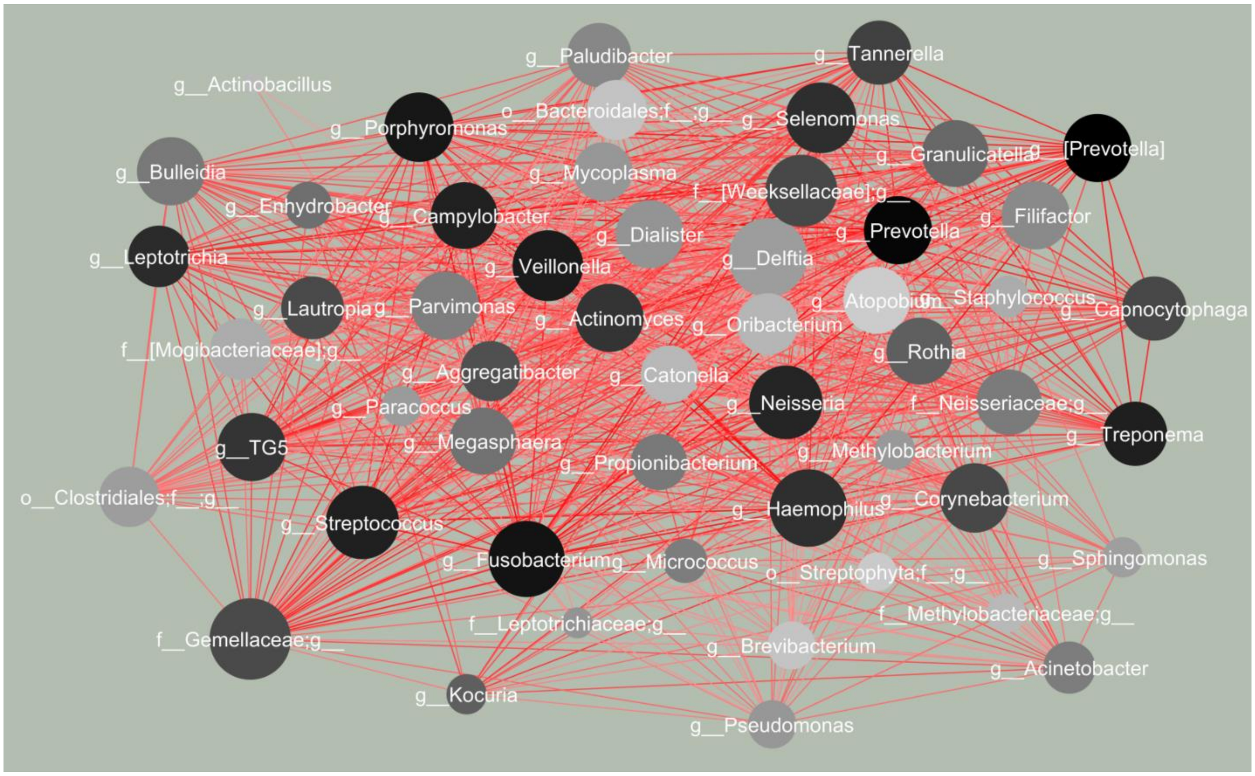


**Figure S11. PC-corr network to investigate the effect of *H. pylori* infection on gastric mucosal microbiota in Parsons *et al.* data.** The PC-corr network was constructed at cut-off 0.5 according to the loadings of PC1, since PCA could significantly (p-value<0.01) separate patients in the normal stomach group (with no evidence of *H. pylori* infection and PPI-untreated, Control) from patients with *H. pylori* gastritis (positive to *H. pylori* infection and not using PPIs, HPGast) (Fig. S9), therefore reflects discriminative network modules related to *H. pylori* infection in gastric mucosa. All the bacteria, that are represented by black nodes, have higher abundance in control group (↑Control), that is their abundance is decreased in the presence of *H. pylori* infection (↓HPGast).

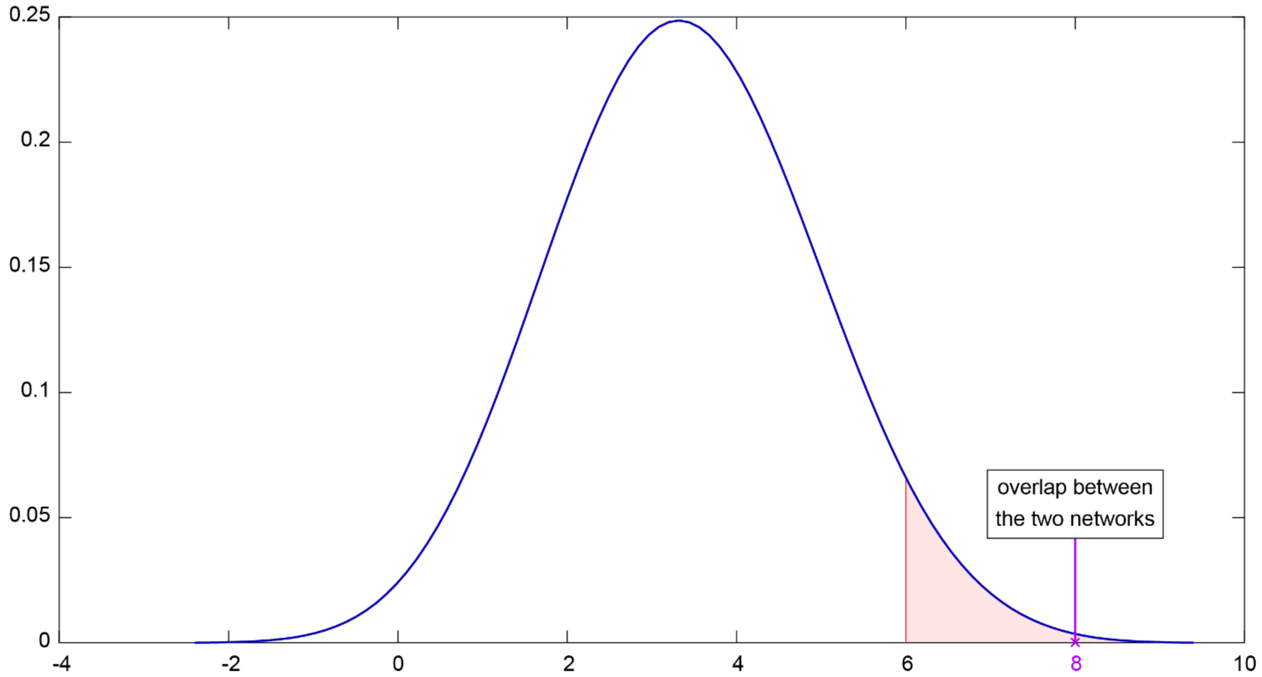


**Figure S12. The overlap between Paroni Sterbini *et al.* and Parsons *et al.* networks, exemplifying the effect of *H. pylori* infection on gastric mucosal microbiota, is statistically different from a random overlap.** To verify that the overlap between the two networks (violet circle in Figure 8) is statistically different from a random overlap, we performed a statistical test based on random resampling of the bacteria in the two networks (repeated 10,000 times). The found intersection (8 bacteria, violet line) is significantly different (p-value=1.00e-04) from a random resampling overlap considering a level of significance of 0.05 (with corresponding critical region in red and critical value denoted with red line), meaning the probability of obtaining at random the same intersection is very low.
